# Phylogenetic analysis of *ABCE* genes across the plant kingdom

**DOI:** 10.1101/2023.09.29.560150

**Authors:** Liina Jakobson, Jelena Mõttus, Jaanus Suurväli, Merike Sõmera, Jemilia Tarassova, Lenne Nigul, Olli-Pekka Smolander, Cecilia Sarmiento

**Author notes:** Authors for Correspondence: Liina Jakobson, Plant Biotechnology Department, Centre of Estonian Rural Research and Knowledge, M. Pilli haru 1, Jõgeva, Estonia; Cecilia Sarmiento, Department of Chemistry and Biotechnology, Tallinn University of Technology, Akadeemia tee 15, 12618, Estonia. Equal contribution (Liina Jakobson, Jelena Mõttus).

## Abstract

ATP-BINDING CASSETTE SUBFAMILY E MEMBER (ABCE) proteins are one of the most conserved proteins across eukaryotes and archaea. Yeast and the vast majority of animals possess a single *ABCE* gene encoding the vital ABCE1 protein. We retrieved *ABCE* gene sequences of 76 plant species from public genome databases and analyzed them with the reference to *Arabidopsis thaliana ABCE2* gene (*AtABCE2*). Over half of the studied plant species possess two or more *ABCE* genes. There can be as many as eight *ABCE* genes in a plant species. This suggest that *ABCE* genes in plants can be classified as a low-copy gene family, rather than a single-copy gene family. Plant ABCE proteins showed overall high sequence conservation, sharing at least 78% of amino acid sequence identity with AtABCE2. The phylogenetic trees of full-length ABCE amino acid and CDS sequences demonstrated that *Brassicaceae* and *Poaceae* families have independently undergone lineage-specific split of the ancestral *ABCE* gene. Other plant species have gained *ABCE* gene copies through more recent duplication events. Deeper analysis of *AtABCE2* and its paralogue *AtABCE1* from 1135 *Arabidopsis thaliana* ecotypes revealed 4 and 35 non-synonymous SNPs, respectively. The lower natural variation in *AtABCE2* compared to *AtABCE1* is in consistence with its crucial role for plant viability. Overall, while the sequence of the ABCE protein family is highly conserved in the plant kingdom, many plants have evolved to have more than one copy of this essential translational factor.

**Significance statement:** In most eukaryotes there is a single ABCE protein, which is involved in many vital processes in cells. However, less is known about ABCEs specifically in plants. Here we show that while the sequence of ABCE proteins is highly conserved in plants, they have evolved to often have multiple copies of this essential translational factor. By studying 76 species from the entire plant kingdom, we observed as many as eight *ABCE* genes being present at a time, although most species have less. Some *ABCE* copies appeared earlier than others and were found in multiple species. Thus, our findings indicate that ABCE genes in plants are not a single-copy gene family and should instead be re-classified as a low-copy gene family.

## Introduction

Members of the ATP-BINDING CASSETTE (ABC) subfamily E (ABCE) belong to the superfamily of ABC proteins, which can be found in all living organisms studied to date and are regarded as highly essential in all eukaryotes. Most ABC proteins function as ATP-dependent membrane transporters. They possess transmembrane domains (TMDs) coupled with nucleotide-binding domains (NBD) otherwise known as ATP-binding cassettes (Andolfo et al. 2015; Navarro-Quiles et al. 2018). ABCE (initially denoted RNASE L INHIBITOR (RLI)) proteins, in contrast, lack TMDs, but still have two NBDs associated with several specific domains and thus are soluble proteins.

In most species the ABCE subfamily is represented by a single member, ABCE1, which is involved in ribosome biogenesis and several stages of translation regulation (Yarunin et al. 2005; Andersen & Leevers 2007; Barthelme et al. 2011; Mancera-Martínez et al. 2017; Navarro-Quiles et al. 2018). In accordance with its fundamental role, ABCE1 expression has been detected in most tissues and developmental stages of the species studied. In addition, loss-of-function of *ABCE1* genes results in a lethal phenotype in all studied species (Du et al. 2003; Zhao et al. 2004; Maeda et al. 2005; Sarmiento et al. 2006; Kougioumoutzi et al. 2013). ABCE1 has been found to participate in translational initiation and termination, however, its most conserved function is in the process linking these two stages of translation – ribosome recycling (Navarro-Quiles et al. 2018). During that process, ABCE1 splits the ribosome through direct interactions with ribosomal subunits and release factors, either after canonical stop codon-dependent termination or after recognition of stalled and vacant ribosomes. The latter is recognized during mRNA surveillance mechanisms such as no-go decay (NGD), non-stop decay (NSD), and non-functional 18S rRNA decay (18S-NRD) (Graille & Séraphin 2012). Furthermore, ABCE1 dissociates the 80S-like complex during maturation of ribosomal subunits (Strunk et al. 2012). The role in ribosome biogenesis is supported by the nuclear accumulation of 40S and 60S ribosome subunits in the absence of ABCE1 (Yarunin et al. 2005; Kispal et al. 2005; Andersen & Leevers 2007). Additionally, it has a key role in RNA silencing in both plants and animals (Zimmerman et al. 2002; Sarmiento et al. 2006; Kärblane et al. 2015). Moreover, we have previously shown that human ABCE1 (HsABCE1) is directly or indirectly involved in histone biosynthesis and DNA replication (Toompuu et al. 2016).

The study of ABCE functions in plants has been mostly limited to the model plants *Arabidopsis thaliana*, *Nicotiana benthamiana*, *Nicotiana tabacum* and *Cardamine hirsuta* (Petersen et al. 2004; Sarmiento et al. 2006; Kougioumoutzi et al. 2013; Mõttus et al. 2021; Navarro-Quiles et al. 2022). In *A. thaliana* there are two genes encoding for paralogous ABCE proteins (AtABCE1 and AtABCE2, also referred to as AtRLI1 and AtRLI2, respectively), which share 80.8% identity (Navarro-Quiles et al. 2022; Mõttus et al. 2021). AtABCE2 is orthologous to HsABCE1 and is ubiquitously expressed in all plant organs (Sarmiento et al. 2006). Recently, AtABCE2 was found to interact with ribosomal proteins and translational factors, confirming its conserved ancestral function in translation that is coupled to general growth and vascular development, likely indirectly via auxin metabolism (Navarro-Quiles et al. 2022). Furthermore, through regulation of translation AtABCE2 is involved in the development of gametophyte and embryo (Yu, et al. 2023). In addition, AtABCE2 has been shown to suppress GFP transgene RNA silencing in heterologous system at the local and at the systemic levels by reducing accumulation of siRNAs (Sarmiento et al. 2006; Kärblane et al. 2015). Mutational analysis of AtABCE2 revealed that the structural requirements for RNA silencing suppression are similar to those needed for ribosome recycling in archaea (Mõttus et al. 2021). This indicates that AtABCE2 might suppress RNA silencing via supporting translation-associated RNA degradation mechanisms. The role of AtABCE1 in *A. thaliana*, which is expressed almost exclusively in generative organs (Navarro-Quiles et al 2022; Yu et al 2023), is yet to be studied.

Silencing of *ABCE* orthologues *(RLIh)* in *N. tabacum* resulted in a single viable transgenic plant exhibiting severe morphological alterations, supporting the important role of ABCE proteins at the whole-organism level. At that time, it remained unclear how many *RLIh* genes there are in tobacco species (Petersen et al. 2004). In *C. hirsuta,* a close relative of *A. thaliana* that has composite leaves, there is only one *ABCE* gene in the genome, named SIMPLE LEAF3 (SIL3, or ChRLI2) (Kougioumoutzi et al. 2013). Hypomorphic mutation Pro177Leu in the NBD1 domain of ChRLI2 affects the determination of leaf shape and regulation of auxin homeostasis (Kougioumoutzi et al. 2013). Interestingly, the expression of *ChRLI2* was not ubiquitous as in *A. thaliana*, but instead it was shown to be expressed in meristematic and vascular tissues of young developing leaves and in leaflet initiation sites (Kougioumoutzi et al. 2013).

It is commonly claimed that most eukaryotes only have one *ABCE* gene (Dermauw & Van Leeuwen 2014). Exceptions to this have been detected in plants such as thale cress, rice, maize, potato and tomato, but also in animals such as catfish, cod and mosquitoes (Braz et al. 2004; Garcia et al. 2004; Verrier et al. 2008; Liu et al. 2013; Pang et al. 2013; Andolfo et al. 2015; Lu et al. 2016). Although some plant species have more than one *ABCE* gene, it is still the smallest and most conserved of all ABC subfamilies (Andolfo et al. 2015).

In this study we aimed to characterize the phylogenetic evolution of *ABCE* genes in plants in order to shed light on the possible functional diversification within ABCE protein family. Here we present the results of an extensive bioinformatics analysis of publicly available sequences for plant *ABCE* genes and corresponding proteins, together with haplotype analysis of *A. thaliana ABCE*s.

## Results

### Variability of plant *ABCE* genes

To gain insight into the diversity of *ABCE* genes in plants, we compiled a selection of *ABCE* genes from 76 different plant species available in public databases. The selection criteria for including in further analysis was high identity (min 78%) of the full-length amino acid sequence to AtABCE2 and the presence of all known essential structural elements of ABCE proteins (Karcher et al. 2005; Barthelme et al. 2007; Nürenberg & Tampé 2013; Nürenberg-Goloub et al. 2020). Truncated or aberrant sequences were discarded from further analysis. Altogether 152 plant *ABCE* genes were included in the study. The selected species represented a wide range of plant groups, including unicellular algae such as *Chlamydomonas reinhardtii* and *Micromonas sp. RCC299*, monocots such as *Zea mays* and *Triticum aestivum*, *Solanum* species such as *Solanum tuberosum* and *N. benthamiana*, and *Brassicaceae* such as *Brassica napus* and *A. thaliana* (Figure 1, Supplementary Table S1). Our analysis revealed that plant species from the phylum *Chlorophyta* (green algae) usually possess only a single *ABCE* gene, except for *Ostreococcus lucimarinus*, which has two genes. In contrast, most of the analyzed species in the *Poaceae* family have at least two *ABCE* genes, and some have as many as eight genes in their genome, as is the case for *T. aestivum*. Another group of plant species with an above-average number of *ABCE* genes is the *Brassicaceae* family. For example, *B. napus* has eight genes, *B. rapa* five genes, and *Capsella rubella* four genes. On the other hand, among *Brassicaceae*, *C. hirsuta* and *Boechera stricta* have only a single *ABCE* gene (Figure 1). Interestingly, only 30 species out of 76 (39.5%) had a single *ABCE* gene. Similarly, there were 34 species (44.7%) possessing two *ABCE* genes (Figure 2A). This shows that *ABCE* genes in plants are not a single-copy gene family and should instead be re-classified as a low-copy gene family.

**Fig. 1.**
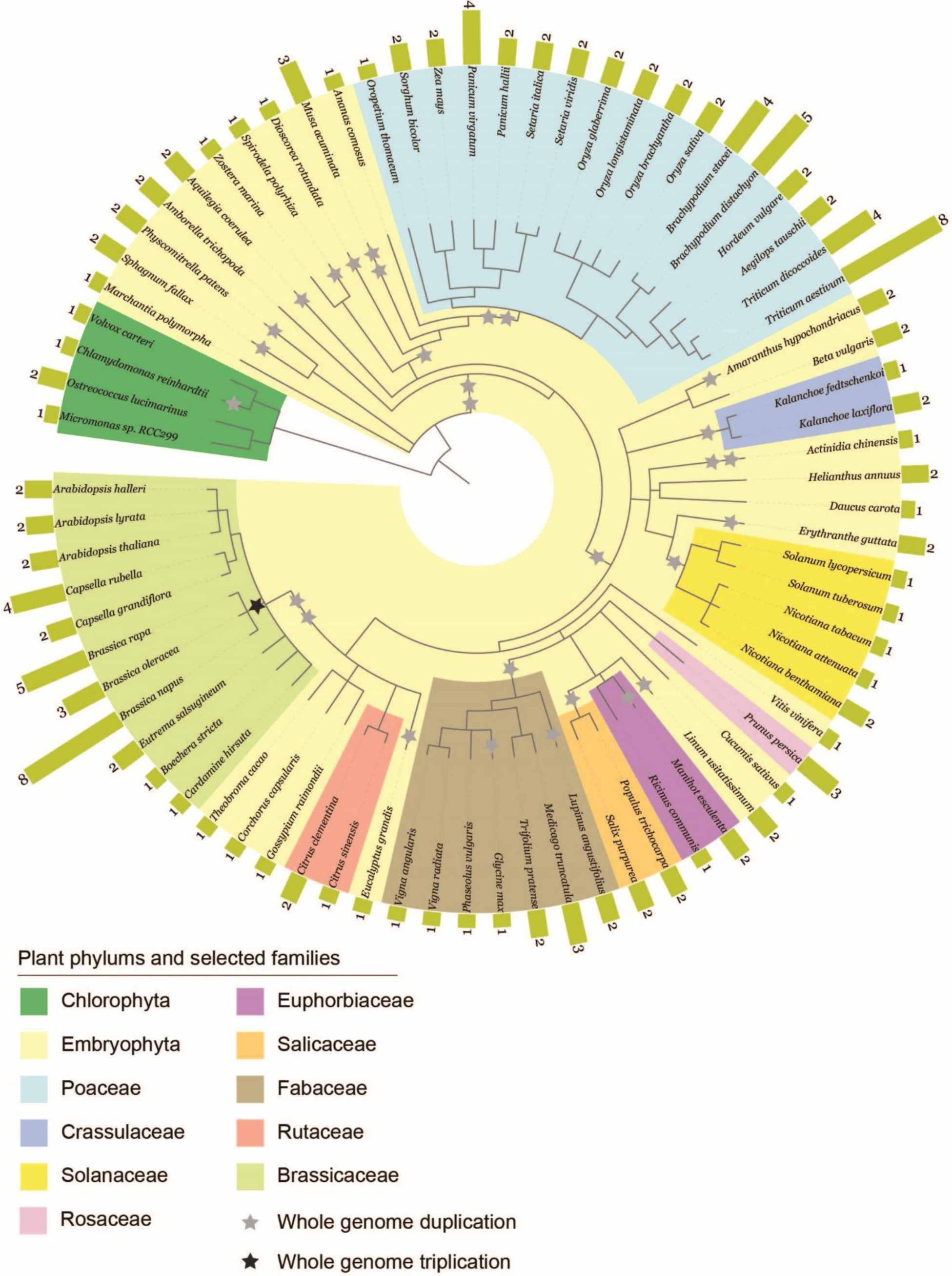
Phylogenetic diagram of 152 *ABCE* genes in plants. The data was compiled from 76 plant species. Whole genome duplications (WGDs) and triplications (WGT) are marked as grey and black stars, respectively.

**Fig. 2.**
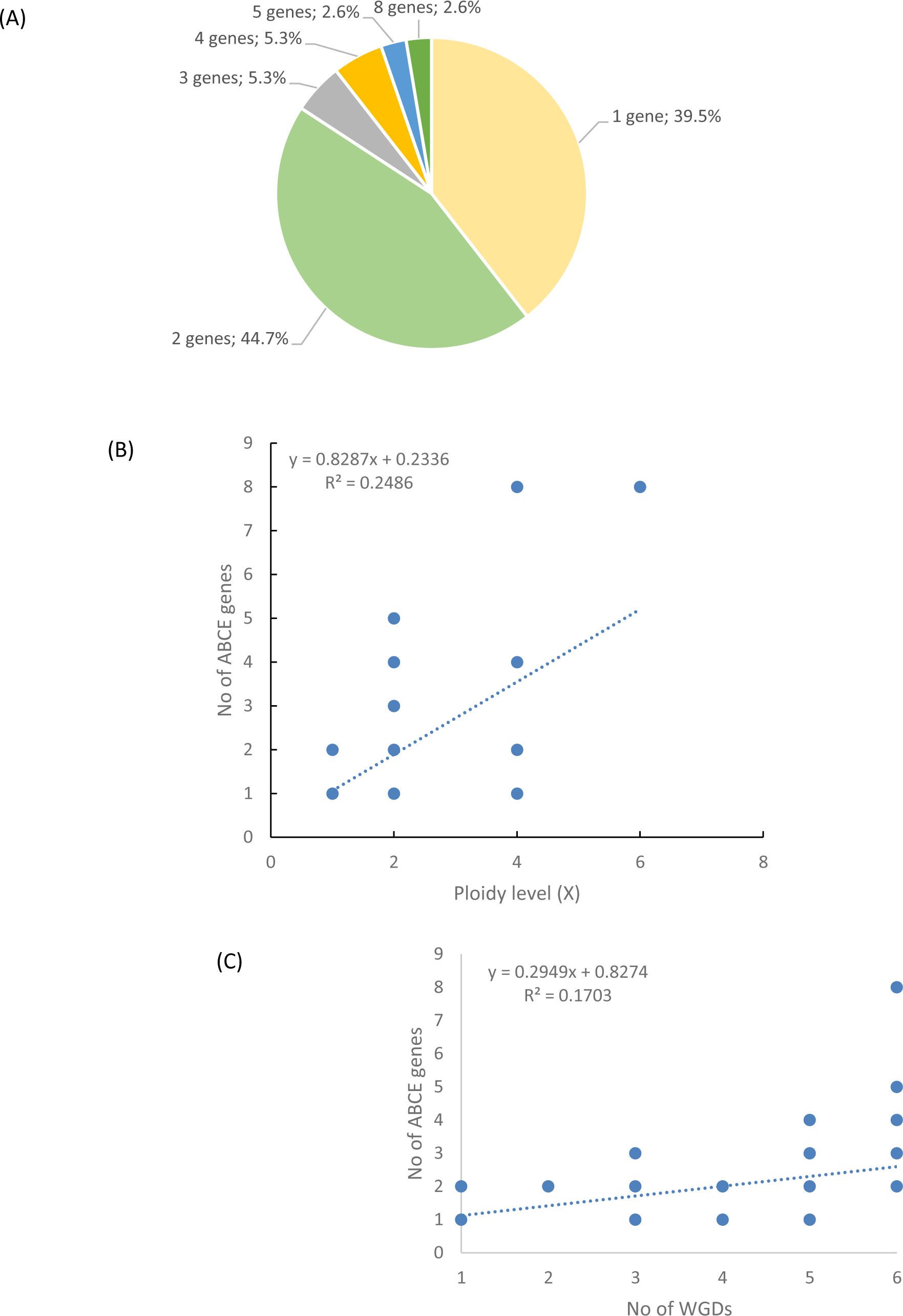
*ABCE* copy number in plants. (A) *ABCE* gene copy number among the studied 76 plant species. (B) The correlation between *ABCE* gene copy number and species ploidy level. (C) The correlation between *ABCE* gene copy number and the number of WGDs of a species.

Next, we analyzed the correlation between ploidy level and the number of *ABCE* genes. Despite clustering of multi-gene-species in the *Poaceae* and *Brassicaceae* families, there was no visual segmentation between the number of *ABCE* genes and phylogenetic origin in other plant families (Figure 1). We found a slight correlation (R=0.25) between ploidy level and the amount of *ABCE* genes per species. However, there were examples of tetraploid species with a single *ABCE* gene (e.g. *S. tuberosum*) and diploids with five *ABCE* genes (e.g. *B. distachyon*, *B. rapa*) (Figure 2B, Supplementary Table S1).

We compared the amount of *ABCE* genes with known whole genome duplications (WGDs) and triplications (WGT) per species. WGD data was based on the data of 53 plant species published by the One Thousand Plant Transcriptomes Initiative (Leebens-Mack et al. 2019). The observation of WGT data in the common ancestor of *Brassica* species was based on the study of Wang and coworkers (Wang et al. 2011). We found a very weak correlation (R=0.17) between the number of *ABCE* genes in a species and the number of WGD events that have taken place in the ancestors of the respective species (Figure 2C). Interestingly, there are examples of species, which have encountered at least five WGDs in their evolutionary history, but still possess only a single *ABCE* gene, for example *Actinidia chinensis* and *Glycine max* (Figure 1, 2C, Supplementary Table S2).

### Plant *ABCE*s are highly conserved

ABCE proteins are composed of four domains: NBD1 and NBD2 forming the ATPase core, bipartite hinge domain that is tightly engaged in twin-NBD cassette arrangement and a unique N-terminal iron-sulphur (FeS) cluster domain (Figure 3A). Additionally, ABCEs embody a helix–loop–helix (HLH) motif in NBD1 that distinguishes it from otherwise superimposable NBD2 (Karcher et al. 2005).

**Fig. 3.**
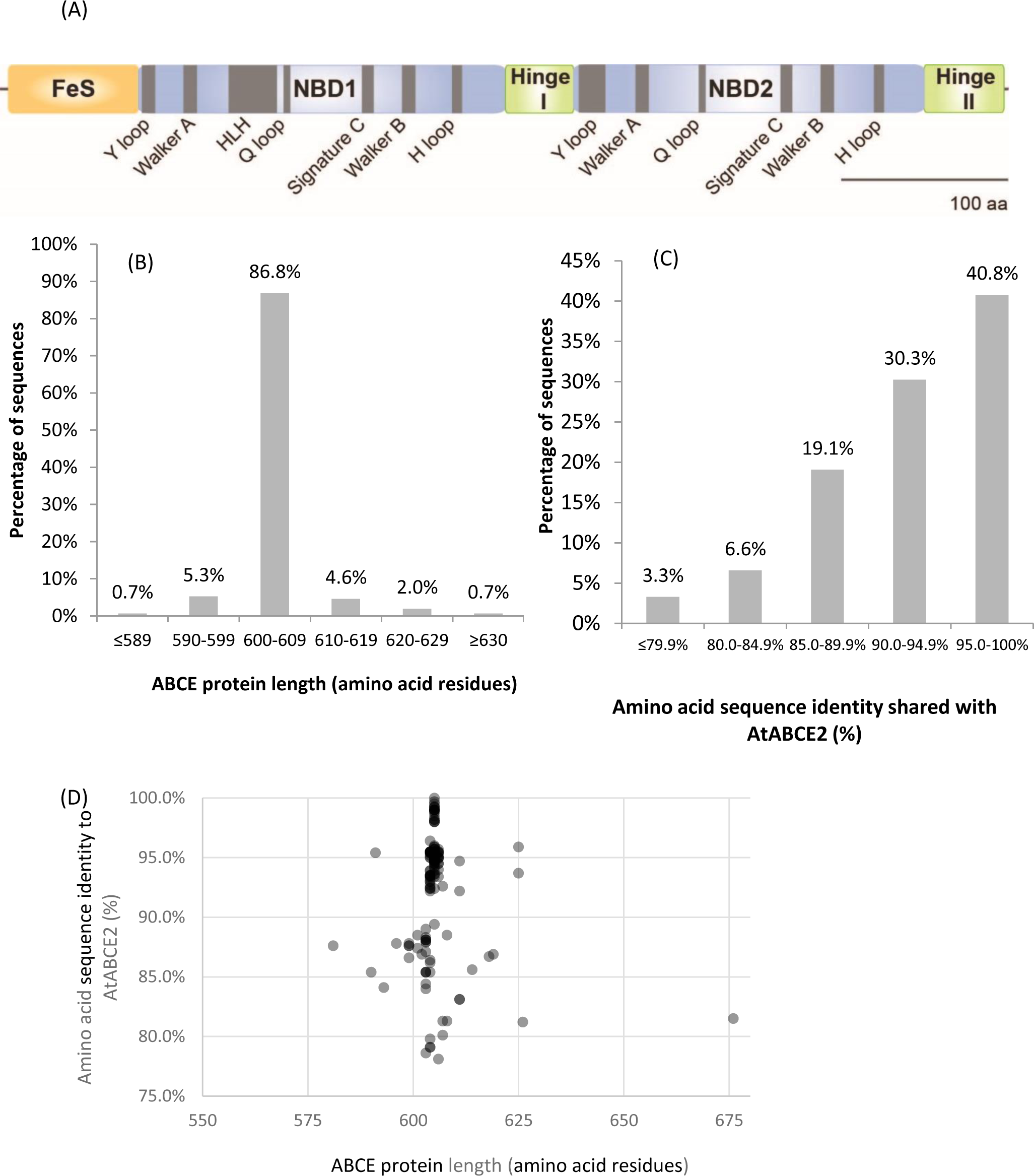
Amino acid sequences of plant ABCEs reveal high level of conservation. (A) Linear protein model of AtABCE2. Grey regions depict highly conserved motifs within AtABCE2. FeS – iron-sulphur cluster domain, NBD1 – nucleotide-binding domain 1, NBD2 – nucleotide-binding domain 2. (B) Histogram of protein sequence lengths of the studied 152 plant ABCEs. (C) Histogram of amino acid sequence identities of the studied 152 plant ABCEs, based on MUSCLE alignment. (D) The correlation between amino acid sequence identity and protein sequence length of the studied 152 plant ABCEs.

As many as 86.8% out of the 152 analyzed gene sequences encode ABCE of canonical protein length (600 – 609 amino acids) (Figure 3B). Along with exceptional conservation within functionally critical motifs, all analyzed ABCE sequences are highly similar to AtABCE2 sharing at least 78% of amino acid sequence identity (Figure 3C, Supplementary Table S3).

We also plotted amino acid sequence length to amino acid sequence identity for the studied 152 ABCE proteins. There was a clear clustering of proteins with the length of 605 amino acids (Figure 3D). Proteins with lower sequence identity did not cluster by protein length (Figure 3D). Interestingly, proteins with more than 90% identity to AtABCE2 could be as short as 591 amino acids and as long as 625 amino acids long (Figure 3D). Hence, despite some variance in amino acid sequence length and sequence identity to AtABCE2, the selection of amino acid sequences analyzed here is uniform and represents well the plant *ABCE* genes.

### Phylogeny of plant *ABCE*s

To understand how ABCE proteins have evolved in the green plant lineage, we constructed a Maximum Likelihood (ML) phylogenetic tree for 152 ABCE protein sequences from 76 species (Figure 4). Among the included species, *Chlorophyta* (green algae) represent the earliest lineage to split off from the rest of the green plants. In an unrooted tree of ABCE amino acid sequences, representatives of *Chlorophyta* formed a separate cluster with high bootstrap support (Supplementary Figure S1a). In a smaller tree with ABCE amino acid sequences from 15 plant species rooted with HsABCE1, the sequences from *Chlorophyta* were clearly split off from all other plants (Supplementary Figure S1b). We therefore reasoned that *Chlorophyta* would be a suitable outgroup for rooting the tree.

**Fig. 4.**
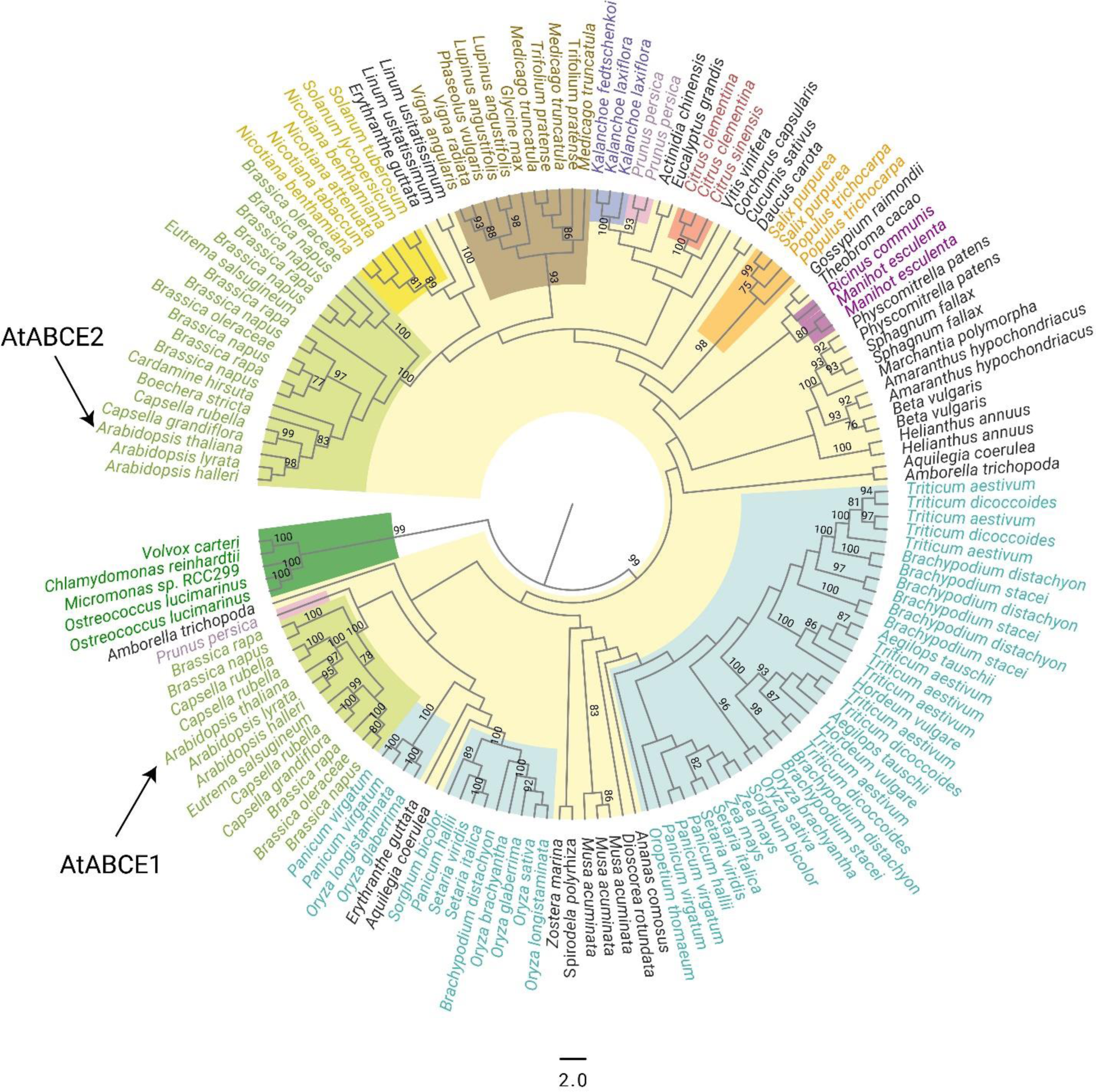
Cladogram of the 152 plant ABCE full-length amino acid sequences. There was a total of 777 positions in the final dataset. The tree was constructed using the Maximum Likelihood method and JTT+G model. Color coding of the selected proteins refers to the affiliation to the bigger plant phyla or families. Bootstrap values over 75% are depicted on the figure. The tree is rooted on *Chlorophyta* phylum.

By examining the tree rooted on *Chlorophyta*, we found that most of the plant ABCE full-length protein sequences cluster well together according to the taxonomic groupings. For instance, ABCEs from *Fabaceae* family members (legumes) form a single cluster, as is the case for *Solanaceae* family (nightshades). Homologous ABCE protein sequences from these families share a very high similarity and likely have the same function (Figure 4). As previously reported for a small selection of *Brassicaceae* (mustard and cabbage family, includes the thale cress *A. thaliana*) (Navarro-Quiles et al. 2022), our analysis shows that many of their members encode distinguishable ABCE1 and ABCE2 proteins (Figure 4). Notably, all *Brassicaceae* species have at least one ABCE2 protein.

In contrast to the well supported clustering of sequences from the same taxonomic groups, relationships between the clusters were not fully resolved and all internal branches of the tree were poorly supported. The topology of the likeliest amino acid sequence-based tree suggested all ABCE proteins from *Poaceae* (grasses) to be more closely related to AtABCE1 than to AtABCE2. This contrasts the notion that AtABCE2 preserves the ancestral function, whereas AtABCE1 is subjected to ongoing subfunctionalization or pseudogenization (Navarro-Quiles et al. 2022). In an attempt to shed further light on the phylogenetic relationships between AtABCE proteins and ABCEs from *Poaceae* family members, we constructed an additional tree including ABCE amino acid sequences from the *Brassicaceae* (32) and *Poaceae* (48) families, together with *Amborella trichopoda* (2), an important reference plant species in evolutionary studies. The green algae *C. reinhardtii* (1) was used to root the tree. In the likeliest tree obtained from this analysis, *Brassicaceae* ABCE1 group appeared as a separate branch, whereas all *Poaceae* ABCE protein sequences clustered together with AtABCE2, a very different result than the one obtained before. This further highlights that the amino acid sequences of ABCE may not be sufficient to fully resolve the phylogeny (Figure 5).

**Fig. 5.**
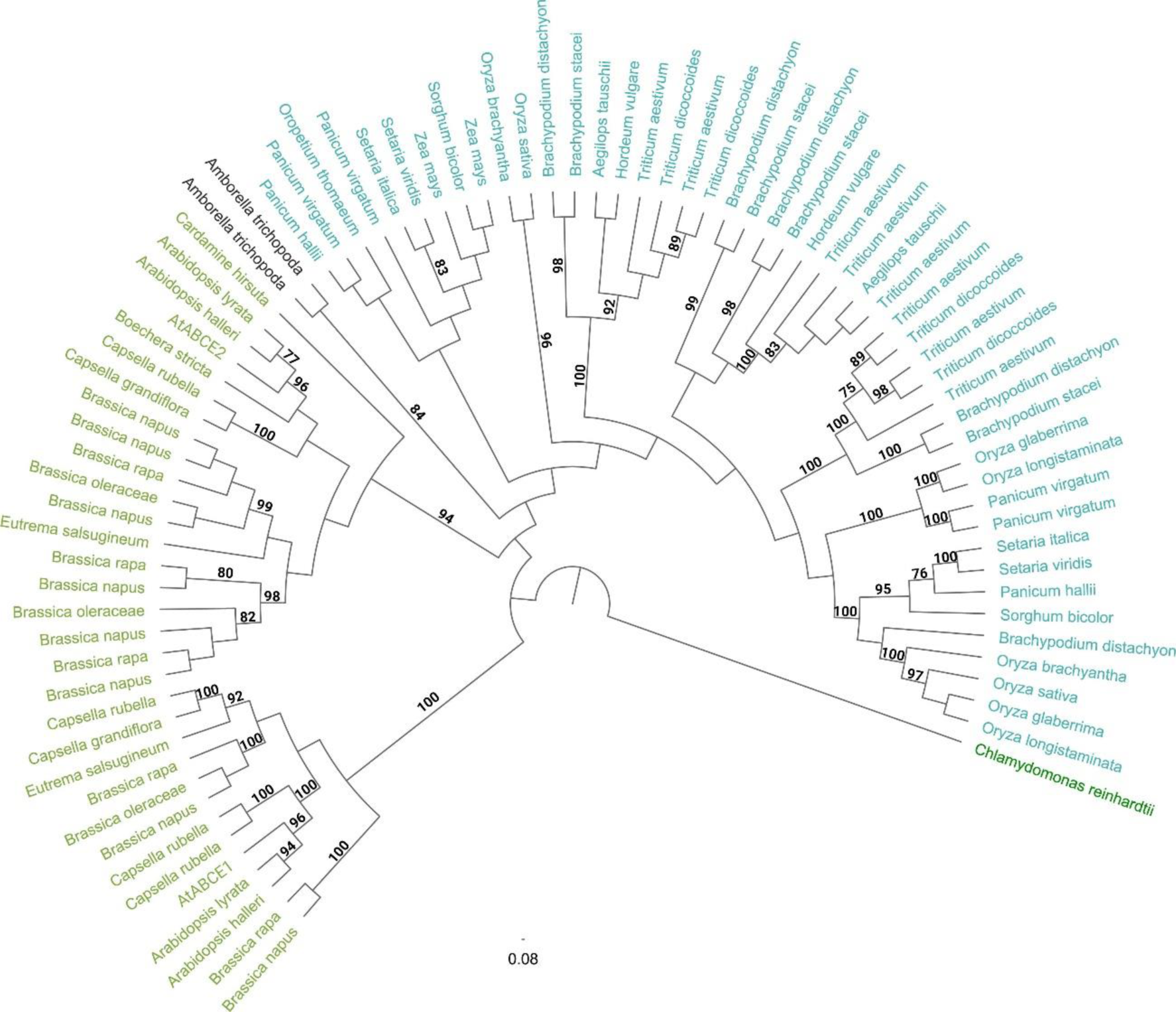
Cladogram of the selected 83 plant ABCE amino acid sequences from the *Brassicaceae* family (32), *Poaceae* family (48) and *A. trichopoda* (2) together with *C. reinhardtii* (1) as the root of the tree. There was a total of 738 positions in the final dataset. The tree was constructed using the Maximum Likelihood method and JTT+G model. Bootstrap values over 75% are depicted on the figure.

We then constructed an additional ML-tree based on nucleotide sequences coding for the ABCE proteins (coding sequence; CDS), rather than the proteins themselves. We used CDS of the same 152 plant ABCE proteins that were used to generate Figure 4, and again used *Chlorophyta* as an outgroup for placing the root (Figure 6). The information provided by nucleotide-level differences allowed additional branches in the tree to be resolved with high bootstrap support values. Here it becomes clear that the split of a single *ABCE* gene into *ABCE1* and *ABCE2* happened in the common ancestor of *Brassicaceae*, whereas the ancestor of *Poaceae* had a similar split independently (Figure 6). Furthermore, it appears that in the common ancestor of the largest *Poaceae* subfamily *Pooideae*, involving important cereals such as barley and wheat, one *ABCE* paralogue went through additional duplication (Figure 6). Although many other plant species also have more than one *ABCE* gene, these originate from more recent duplication events that are usually not shared with other analyzed species. The placement of major clusters in relation to each other generally correlates with the species tree (Figure 6).

**Fig. 6.**
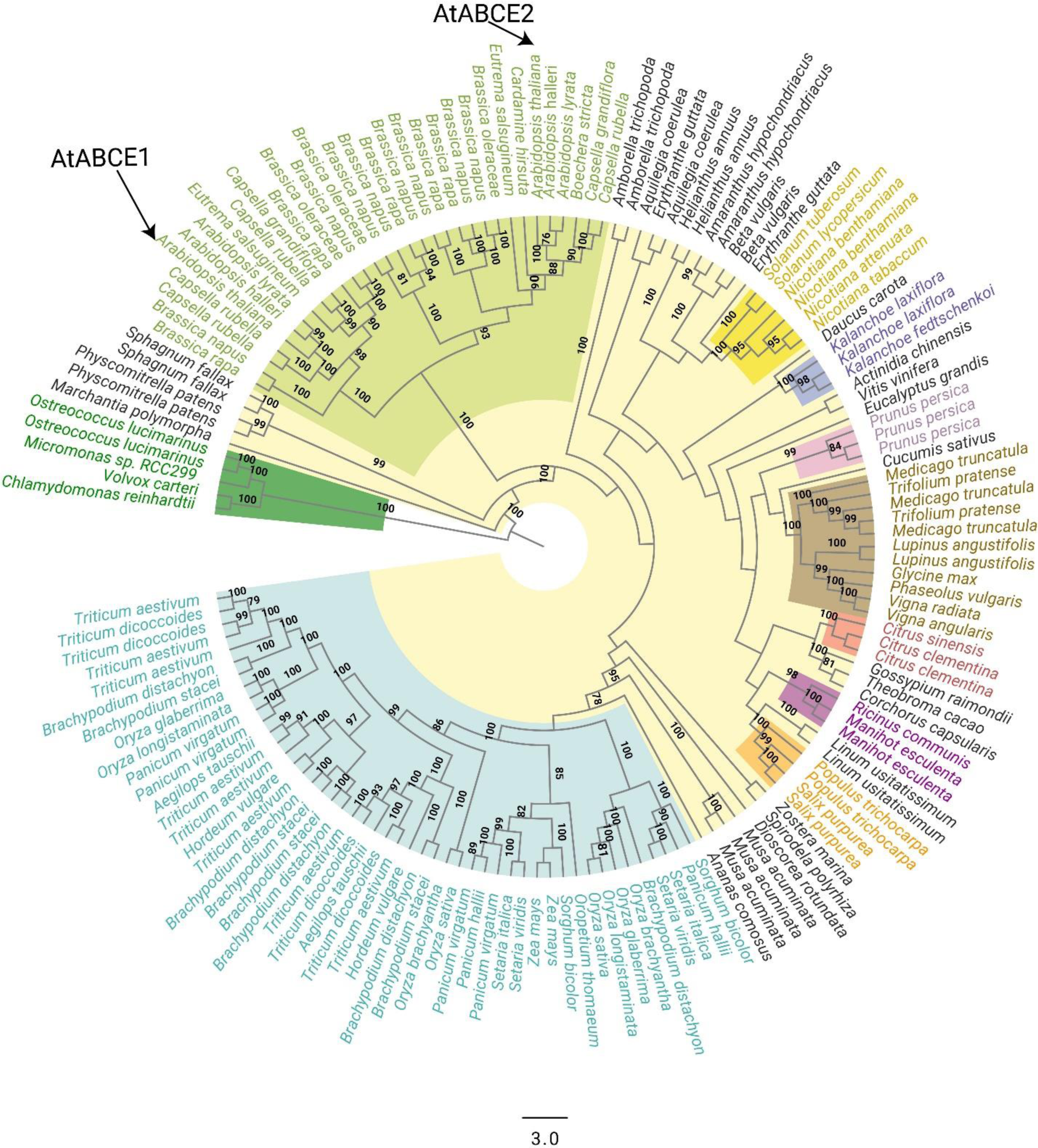
Cladogram of the 152 plant *ABCE* full-length CDS sequences. There was a total of 1810 positions in the final dataset. The tree was constructed using the Maximum Likelihood method and JTT+G model. Color coding of the selected proteins refers to the affiliation to the bigger plant phyla or families. Bootstrap values over 75% are depicted on the figure.

In addition, we separately analyzed the three main domains of ABCE proteins, FeS cluster domain, NBD1, and NBD2, by realigning the corresponding sequences and constructing a ML-tree for each (Supplementary Figure S2, S3, S4). The topology of the resulting trees was different for each domain, but none of those were well supported by bootstrap analysis.

### Natural variation of Arabidopsis *AtABCE1* and *AtABCE2* genes

Natural variation among *A. thaliana* ecotypes has been well documented by the 1001 Genomes Project (Weigel & Mott 2009). We analyzed the *ABCE* gene sequences of all 1135 *A. thaliana* ecotypes reported in that project and found 35 and 4 non-synonymous SNPs in *AtABCE1* and *AtABCE2*, respectively (Supplementary Table S4). Only four non-synonymous SNPs in *AtABCE2* indicate a low degree of natural variation, which is consistent with its fundamental, conserved role in growth and development. On the other hand, 35 non-synonymous SNPs annotated for *AtABCE1* show relatively higher natural variation. From these SNPs, we selected 18 that cause amino acid substitutions at conserved and important positions or that were present in combination with other SNPs of interest. Therefore, 21 ecotypes were included in the further study and resequencing, together with Col-0 (Supplementary Table S4; Table 1). Two SNPs causing amino acid substitution could not be detected while resequencing (Pro399Thr in Grivo-1 and Gly182Ser in IP-Cot-0). Instead, one SNP previously undocumented in the 1001 Genomes Project database (Leu253Phe in Grivo-1) was identified. Figure 7A shows the positions of the amino acid substitutions caused by the 17 SNPs sequenced in the *AtABCE1* gene. In our resequencing analysis, the most frequent SNPs in *AtABCE1* caused the changes His561Leu and Ala441Thr (Table 1). Noteworthily, histidine at the position 561 seems to be characteristic to Col-0, as all the other analyzed ecotypes had leucine at this position. Next, we performed haplotype analysis with CDS sequences on the PopART platform and found that the most conserved sequence of *AtABCE1* is most probably the one identical to Ei-2, Kia1, Pra-6, IP-Car-1 and Can-0 (Figure 7B). Eight SNPs out of 17 appear as single SNPs in the *AtABCE1* of Kly4, Toufl-1, Col-0, IP-Ezc-2, IP-Vis-0, IP-Moz-0, IP-Hoy-0, IP-Loz-0, IP-Cot-0 and Lebja-1 (Figure 7B). Interestingly, a substitution of Ala441Thr can appear both as the consequence of a single SNP in Leska-1-44 and together with other SNPs such as in Cvi-0 or Qar-8a (Figure 7B). Some amino acid changes, like Pro129Gln, Ala549Gly and His561Leu, are always grouped (Table 1). Pro129Gln appears only together with at least two other SNPs, e.g., in Grivo-1 or Qar-8a (Figure 7B).

**Fig. 7.**
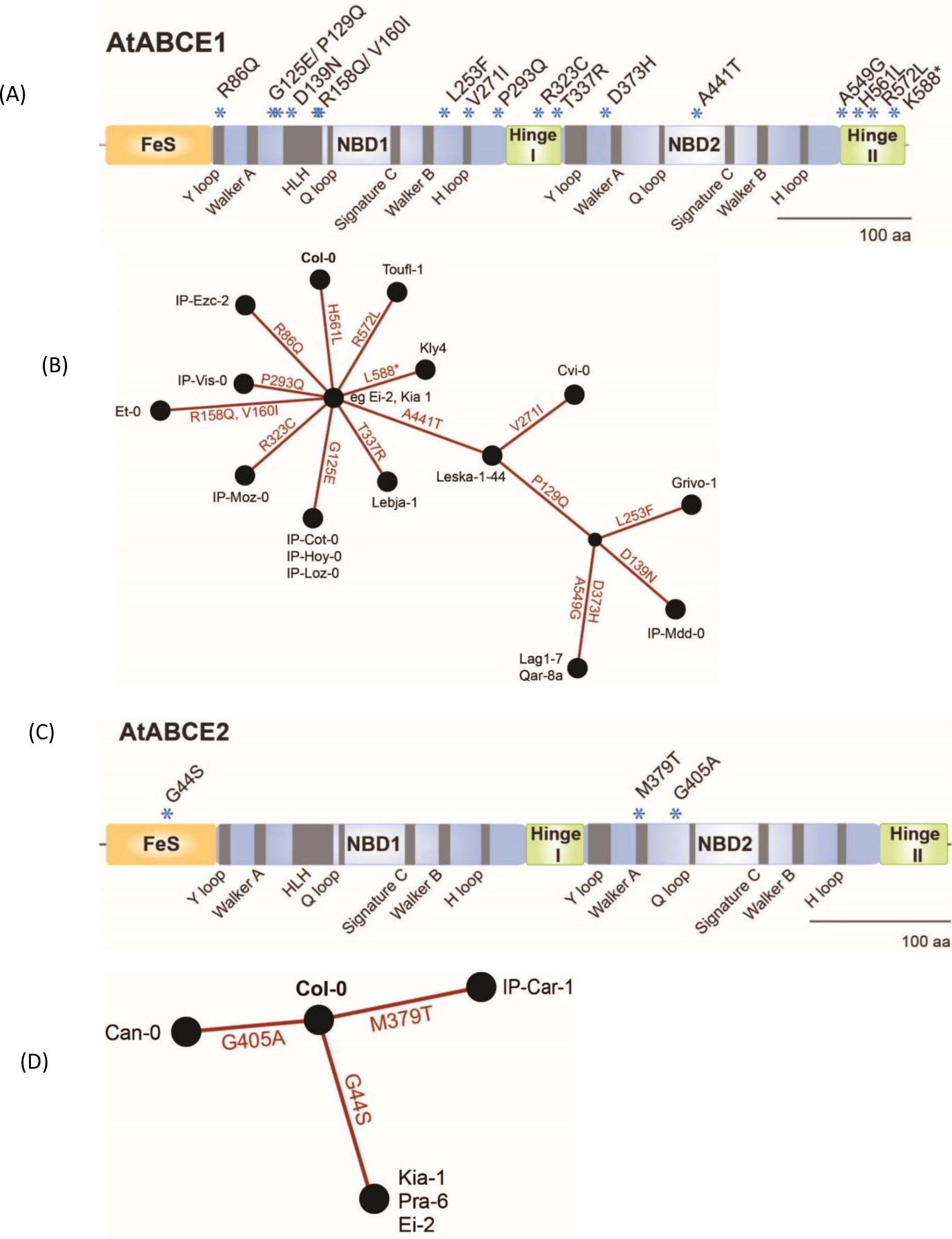
Analysis of non-synonymous SNPs in *AtABCE1* and *AtABCE2*. (A) Linear protein model of AtABCE1. Asterisks depict amino acid substitutions due to SNPs verified in different ecotypes. (B) Haplotype map of AtABCE1 detected among 22 *Arabidopsis* ecotypes. Branch length represents the number of mutations between sequences. For pairs of haplotypes whose distances on the tree are longer than the distances between the sequences, edges are added to shorten the distance. (C) Linear protein model of AtABCE2. Asterisks depict amino acid substitutions due to SNPs verified in different ecotypes. (D) Haplotype map of AtABCE2 detected among six *Arabidopsis* ecotypes. Branch length represents the number of mutations between sequences. For pairs of haplotypes whose distances on the tree are longer than the distances between the sequences, edges are added to shorten the distance.

**Table 1.** Non-synonymous SNPs found in *AtABCE1* and *AtABCE2* among 21 *A. thaliana* ecotypes. All SNPs were verified by Sanger sequencing and whole-genome sequencing published in the 1001 Genomes project (Weigel & Mott 2009; Cao et al. 2011). SNP locations were numbered according to the position in the cDNA sequence starting from ATG. Change in amino acid sequence corresponding to the SNP is presented. Orange color depicts SNPs that are not present in 1001 Genomes Project data but verified by Sanger sequencing within this study. Green color shows SNPs positioned in conserved arginine residues.

| No | Ecotype | SNPs in *AtABCE1* cDNA | Amino acid change in *AtABCE1* | SNPs in *AtABCE2* cDNA | Amino acid change in *AtABCE2* | Origin |
| --- | --- | --- | --- | --- | --- | --- |
| 1 | IP-Ezc-2 | 257G>A<br>1682A>T | Arg86Gln<br>His561Leu | None | None | Spain |
| 2 | IP-Cot-0 | 374G>A<br>1682A>T | Gly125Glu<br>His561Leu | None | None | Spain |
| 3 | IP-Hoy-0 | 374G>A<br>1682A>T | Gly125Glu<br>His561Leu | None | None | Spain |
| 4 | IP-Loz-0 | 374G>A<br>1682A>T | Gly125Glu<br>His561Leu | None | None | Spain |
| 5 | IP-Vis-0 | 878C>A<br>1682A>T | Pro293Gln<br>His561Leu | None | None | Spain |
| 6 | IP-Moz-0 | 967C>T<br>1682A>T | Arg323Cys<br>His561Leu | None | None | Spain |
| 7 | Lebja-1 | 1010C>G<br>1682A>T | Thr337Arg<br>His561Leu | None | None | Russia |
| 8 | Leska-1-44 | 1321G>A<br>1682A>T | Ala441Thr<br>His561Leu | None | None | Bulgaria |
| 9 | Toufl-1 | 1682A>T<br>1715G>T | His561Leu<br>Arg572Leu | None | None | Morocco |
| 10 | Kly4 | 1682A>T<br>1762A>T | His561Leu<br>Lys588STOP | None | None | Russia |
| 11 | Et-0 | 473G>A<br>478G>A<br>1682A>T | Arg158Gln<br>Val160Ile<br>His561Leu | None | None | France |
| 12 | Cvi-0 | 811G>A<br>1321G>A<br>1682A>T | Val271Ile<br>Ala441Thr<br>His561Leu | None | None | Cape Verde |
| 13 | IP-Mdd-0 | 386C>A<br>415G>A<br>1321G>A<br>1682A>T | Pro129Gln<br>Asp139Asn<br>Ala441Thr<br>His561Leu | None | None | Spain |
| 14 | Grivo-1 | 386C>A<br>757C>T<br>1321G>A<br>1682A>T | Pro129Gln<br>Leu253Phe<br>Ala441Thr<br>His561Leu | None | None | Bulgaria |
| 15 | Lag1-7 | 386C>A<br>1117G>C<br>1321G>A<br>1646C>G<br>1682A>T | Pro129Gln<br>Asp373His<br>Ala441Thr<br>Ala549Gly<br>His561Leu | None | None | Georgia |
| 16 | Qar-8a | 386C>A<br>1117G>C<br>1321G>A<br>1646C>G<br>1682A>T | Pro129Gln<br>Asp373His<br>Ala441Thr<br>Ala549Gly<br>His561Leu | None | None | Lebanon |
| 17 | Ei-2 | 1682A>T | His561Leu | 130G>A | Gly44Ser | Germany |
| 18 | Kia 1 | 1682A>T | His561Leu | 130G>A | Gly44Ser | Sweden |
| 19 | Pra-6 | 1682A>T | His561Leu | 130G>A | Gly44Ser | Spain |
| 20 | IP-Car-1 | 1682A>T | His561Leu | 1136T>C | Met379Thr | Spain |
| 21 | Can-0 | 1682A>T | His561Leu | 1214G>C | Gly405Ala | Spain |

In *AtABCE2*, we found only three amino acid changes (Gly44Ser, Met379Thr and Gly405Ala) due to non-synonymous SNPs, which were located in FeS cluster domain and NBD2 at not conserved positions (Figure 7C; Supplementary Table S4). The SNPs were verified in five different ecotypes (Table 1). The haplotype map of *AtABCE2* SNPs shows that Col-0 has the most conserved sequence and three different SNPs root from it (Figure 7D).

In this study we could not find any correlation between the presence of non-synonymous SNPs in *AtABCE* genes and the geographical origin of the ecotype (Table 1). Visual rosette phenotype of the studied ecotypes matched with characterization available in the public databases (Supplementary Figure S5).

In the case of 18 out of 21 ecotypes, all SNPs were confirmed as reported earlier. For three ecotypes only part of the SNPs was validated: Leska-1-44 did not exhibit Pro129Gln, IP-Cot-0 did not exhibit Gly182Ser and Grivo-1 did not exhibit Pro399Thr amino acid changes in AtABCE1 (Supplementary Table S4; Table 1). More interestingly, we verified Gly125Glu in IP-Cot-0 and Pro129Gln, Leu253Phe, Ala441Thr and His561Leu in Grivo-1. Leu253Phe had not been annotated in any *A. thaliana* ecotype in the 1001 Genomes Project database (Table 1).

Surprisingly, we noticed some SNPs affecting highly conserved amino acid residues in AtABCE1. These include arginine residues from R cluster of Hinge domains (Arg323Cys and Arg572Leu of IP-Moz-0 and Toufl-1, respectively), and Arg86Gln of IP-Ezc-2 ecotype that locates to the Y-loop I (Figure 7 A; Table 1). According to the 1001 Genomes Project database the latter SNP is present in 19 *A. thaliana* ecotypes (Supplementary table S4).

Although, the length of AtABCE proteins in *A. thaliana* ecotypes is very conserved, in a single ecotype, namely Kly-4, we found a SNP in *AtABCE1* causing premature stop codon that makes the protein 14 amino acid residues shorter. Despite this deletion, the cluster of arginine residues remains intact in Hinge II subdomain (Supplementary Figure S5).

## Discussion

The availability of high-quality plant genome sequences is growing day by day, which creates a completely new and underexploited repository. It has been recognized that the plant genome evolution has been very complex, including polyploidy, periods of rapid speciation and extinction (Leebens-Mack et al. 2019). Interestingly, massive expansions of gene families took place before the origins of green plants, land plants and vascular plants (Leebens-Mack et al. 2019). Whole genome duplications (WGDs) that have occurred at least 244 times throughout the evolution of plants and ferns (Leebens-Mack et al. 2019) increase ploidy of genomes and largely impact gene family size variation within different lineages. Apart from autopolyploidy, which results from intraspecies WGD events, there are also allopolyploid species, which originate from interspecies hybrids and render gene evolution tracking challenging.

### How many *ABCE* genes plants have and need?

As was previously mentioned, in most animal and in yeast species the *ABCE* gene family is represented by a single gene that encodes the vital ABCE1 protein. In plant kingdom, *ABCE* gene family size across different lineages is more variable. Based on the data from the public databases and our analysis we were able to reconfirm the same number of *ABCE* genes for a selection of plant species. For example there is a single gene in *C. hirsuta* (Kougioumoutzi et al. 2013), in *C. reinhardtii* (Li et al. 2022), in *Citrus sinesis* and in *Theobroma cacao* (Navarro-Quiles et al. 2022). Similarly to previous studies, we reverified two genes in *Zea mays* (Pang et al. 2013), in tomato (Ofori et al. 2018), in rice, and in *Populus trichocarpa* (Navarro-Quiles et al. 2022). The same was true for five *ABCE* genes from *Brassica rapa* (Navarro-Quiles et al. 2022). Intriguingly, Zhang and colleagues found three *ABCE* genes in barley, whereas our study identified only two sequences with all canonical subunits (604 and 611 amino acids long) (Zhang et al. 2020). Moreover, for *Capsella rubella* we identified four *ABCE* genes as opposed to two sequences analyzed earlier (Navarro-Quiles et al. 2022). Taken together, we found that plant *ABCE* genes do not comprise a single-copy gene family, but rather should be classified as a low-copy gene family.

In terms of evolutionary perspective, the duplicates of genes resulting from WGD are often subject to relaxed selection, meaning rapid mutation resulting in loss of function (Qiao et al. 2019). In this study we did not notice significant correlation between ABCE gene family size and WGD events occurred in a lineage, probably due to the elimination of defunctionalized *ABCE* duplicates in many species during evolution. For example, in *Glycine max* and *Actinidia chinensis* after five documented WGD events they retained a single *ABCE2* gene. Interestingly, the overexpression of ABCE1 in yeast causes growth inhibition (Dong et al. 2004), meaning that the amount of ABCE present – and therefore probably also the number of hypothetical redundant genes – is critical for the well-functioning of translation, a crucial process.

However, when higher expression of a particular gene is beneficial, its duplicate might be retained in the genome. This could be the case for the two ABCE paralogues in maize that are located close to each in our phylogenetic analysis (Figure 4) and share the same expression pattern profiles (Pang et al. 2013). Alternatively, as a result of faster evolution, gene duplicates may obtain novel functions or specialized expression patterns (Prince & Pickett 2002). In Arabidopsis, AtABCE1 and AtABCE2 exhibit partial functional redundancy. In contrast to AtABCE2, which is ubiquitously expressed, AtABCE1 is mostly present in generative organs and at relatively low levels (Yu et al. 2023; Klepikova et al. 2016; Navarro-Quiles et al. 2022). This could mean an ongoing process of pseudogenization or subfunctionalization, where the paralogues acquire specific roles. There is a growing evidence regarding ribosomal heterogeneity and the existence of specialized cell-type-specific ribosomes (Xue & Barna 2012; Barna et al. 2022), suggesting that AtABCE1 is involved in the regulation of translation in generative tissues. Paralogous *ABCE* genes in plants may serve to provide specificity in fine-tuning translation and controlling cellular translatome (Gerst 2018).

Our initial observations based on a simple correlation analysis revealed a weak correlation between ploidy level and *ABCE* gene number. In the future, the determinants of *ABCE* copy number can be further elucidated by including more species from diverse lineages and using statistical modelling that takes phylogenetic structuring of the data also into account. These models could also potentially include other information from the species that was not used for the present study, such as whether the species is annual or perennial, their preferred mode of reproduction, or what kind of environments do they grow in.

### In plants *ABCE* genes are prone to duplicate

In this study we analyzed 152 ABCE sequences from 76 plant species. This included the most well studied plant ABCE gene – AtABCE2, which is thought to preserve the ancestral functions of ABCE proteins (Navarro-Quiles et al. 2022). Phylogenetic tree of full-length ABCE protein sequences confirmed previously reported clustering into ABCE2 and ABCE1 groups for *Brassicaceae* (Navarro-Quiles et al. 2022) but was unable to resolve the relationship between ABCEs from different families. The lack of phylogenetic signal can at least partially be explained by the strong conservation of plant ABCE proteins, each sharing at least 78% of sequence identity with AtABCE2 (Supplementary Table S3). However, using the nucleotide sequences coding for ABCE proteins allowed to better illustrate the evolution and clustering of plant *ABCE* genes. The results suggest that *Brassicaceae* and *Poaceae* families have undergone independent lineage-specific splits of the ancestral *ABCE* gene. *Pooideae*, the largest *Poaceae* subfamily that includes barley and wheat, appears to have had further duplication events and its members have additional *ABCE* genes. In addition to *Brassicaceae* and *Poaceae*, many other plant taxa have also gained *ABCE* gene copies, most likely as a result of more recent duplications. We can therefore postulate that in contrast to species which possess a single *ABCE* gene and are sensitive to copy number changes (Dong et al. 2004), plants have evolved to benefit from higher numbers of this essential translational factor. Gene copies may arise from different events including WGD, tandem- and transposon-related duplications, but the precise source of *ABCE* subfamily expansion in plants remains to be investigated.

### Natural variation of *A. thaliana* ABCEs

Usually, essential genes are subject to strong evolutionary pressure and thus, non-synonymous SNPs in conserved regions of gene sequences are rare (Pang et al. 2016; Castle 2011). In *AtABCE1* we found three SNPs that could potentially impact the protein’s function (Table1; Figure 7A). SNPs causing the substitutions Arg323Cys (in IP-Moz-0) and Arg572Leu (in Toufl-1) located at Hinge domain I and II could be of importance, since these domains are essential for NBD-twin cassette assembly in the case of ABCE1 in other organisms (Karcher et al. 2005). In addition, AtABCE1 of IP-Ezc-2 ecotype contains an amino acid substitution at position Arg86Gln, which is exceptionally conserved across archaea and eukaryotes and locates to Y-loop I. In the context of Y-loop with consensus sequence H**<u>R</u>**YGVNAF, the arginine residue has been shown to mediate interaction between FeS cluster domain and NBD1 in the sole *ABCE1* gene of *Pyrococcus abyssi* (Karcher et al. 2008).

The His561Leu amino acid change was reported to be present in 997 out of 1135 *A. thaliana* ecotypes (Supplementary Table S4), which suggest that histidine at this position of *AtABCE1* might be characteristic only to a small subset of ecotypes including Col-0. Thus, it seems that leucine is the most conserved residue at position 561 in *AtABCE1* among *Arabidopsis* ecotypes.

As expected, in *AtABCE2* gene, known to be essential for the viability of an organism (Navarro-Quiles et al. 2022; Yu et al. 2023) only four non-synonymous SNPs residing in non-conserved regions were found among 1135 *A. thaliana* ecotypes (Supplementary Table S4). Importantly, the *AtABCE2* gene seems to be hard to mutate, since up to now there is no T-DNA homozygous line available and only one viable, hypomorphic allele has been recently isolated after ethyl methanesulfonate mutagenesis (Navarro-Quiles et al., 2022). Interestingly, the only non-synonymous SNP present in more than one ecotype in the case of *AtABCE2* is leading to Gly44Ser substitution in FeS domain. According to the 1001 Genomes Project database this mutation is present in 54 ecotypes, three of them were confirmed in the current study (Table 1; Figure 7; Supplementary Table S4).

Taken together, this study has shown the surprisingly high number of *ABCE* genes among the plant kingdom. We hypothesize that plants have developed a number of specialized ABCEs with more specific functions compared to species carrying a single copy of *ABCE* gene such as humans, fruit fly or yeast.

## Methods

### Phylogenetic diagram of studied plants

The phylogenetic diagram of the studied plants species together with the bar chart of *ABCE* gene number was created based on NCBI taxonomy with phyloT and visualized with iTOL (Letunic & Bork 2007, 2019; phylot.biobyte.de).

### Genome and proteome data acquisition

ABCE sequence data for 55 species was downloaded from the online resource Phytozome portal https://phytozome.jgi.doe.gov/ (Goodstein et al. 2012). ABCE sequence data for additional 18 species was downloaded from Ensembl Plants (Howe et al. 2020). The genome data for *C. hirsuta* was accessed at http://bioinfo.mpipz.mpg.de/blast/ (Gan et al. 2016). Genome data for *N. tabacum* and *N. benthamiana* was downloaded from Sol Genomics Network http://solgenomics.net (Edwards et al. 2017; Kourelis et al. 2019). We downloaded genomic, CDS and translated amino acid sequences for each plant *ABCE* gene used in the study.

The length of amino acid sequences was calculated with SeqinR package (version 3.6.1) in R 4.0.2 (Charif & Lobry 2007). In order to calculate their similarities to AtABCE2, all 152 sequences were aligned using the online interface of MUSCLE with default Pearson/FASTA parameters provided by the European Bioinformatics Institute (EBI) (Madeira et al. 2019). Thereafter the percent identity scores were calculated with MUSCLE algorithm for aligned sequences in R 4.0.2 package Bio3D version 2.4-1 (Grant et al. 2021; Edgar 2004).

Next, sequences aligned to AtABCE2 were inspected for the general protein structure, that is the presence and correct order of the domains, including FeS cluster domain, NBD1, NBD2 and bipartite Hinge domain (Supplementary Figure S6). Sequences lacking critical motifs within these domains (Karcher et al. 2005; Barthelme et al. 2007; Nürenberg & Tampé 2013; Nürenberg-Goloub et al. 2020) were filtered out from the analysis.

### Evolutionary analysis by Maximum Likelihood method

All amino acid sequences were analyzed with the MEGA software package (versions 10.2.2 and 11.0.13) (Kumar et al. 2018) as described below. Sequences were aligned with default parameters of the MUSCLE algorithm. Phylogenetic relationships were inferred using the Maximum Likelihood method and JTT matrix-based model (Jones et al. 1992), selected for each data set based on the lowest BIC scores (Bayesian Information Criterion). A discrete gamma distribution was used to model evolutionary rate differences among sites. Bootstrap analysis was performed with 500 replicates. The trees with the highest log likelihood were published for each analysis.

CDS sequences were aligned with the default parameters of MAFFT version 7.4.9.0 (Katoh & Standley 2013). Sites in the CDS alignment that contained gaps for more than 10% of the sequences were removed with trimAl version 1.4.rev22 (Capella-Gutiérrez et al. 2009), resulting in 1,810 retained positions. RAxML version 8.2.12 (Stamatakis 2014) was used to create a maximum likelihood phylogeny with the GTR model of nucleotide substitution, gamma model of rate heterogeneity and 500 bootstrap replicates.

### Data acquisition for 1135 Arabidopsis ecotypes

Data (SNPs and indels) available for 1135 *A. thaliana* strains was downloaded from the 1001 Genomes Project depository (Weigel & Mott 2009).

### Reconfirming the haplotypes of Arabidopsis ecotypes

Seeds of the selected 21 Arabidopsis ecotypes were acquired from the Nottingham Arabidopsis Stock Centre (NASC). Both *AtABCE1* and *AtABCE2* full coding sequences were PCR-amplified and sequenced by Sanger sequencing for ecotypes Can-0, Ei-2, IP-Car-1, Kia1 and Pra-6. For the other 15 ecotypes only *AtABCE1* full coding sequences was sequenced. Primer pairs used for the PCRs are shown in the Supplementary Table S5. For all the PCR reactions touchdown PCR method with the following conditions was used: 95 °C for 15 minutes; 13 cycles of at 95 °C for 15 seconds, at the gradually decreasing temperature from 60 °C to 54 °C (the temperature drops by 0.5 °C per cycle) for 30 seconds, at 72°C for 70 seconds; 15 cycles of 95°C 15 seconds, 54°C 30 seconds, 72°C 70 seconds and the final extension at 72 °C for 10 minutes. Amplified DNA fragments were purified from the agarose gel using GeneJET Gel Extraction Kit (Thermo Scientific) according to the manufacturer’s instructions. Thereafter the purified DNA fragments were Sanger sequenced and aligned respectively to *AtABCE1* or *AtABCE2*. The final results were based on the sequencing of at least two plants for each ecotype. Columbia (Col-0) ecotype was used as a reference.

### Generating haplotype map with PopART

For the haplotype analysis, first an alignment file with CDS sequences was created in Nexus format. Thereafter the multiple sequence alignment was analyzed with PopART version 1.7 (Population Analysis with Reticulate Trees) (Leigh & Bryant 2015). The network was constructed with Median Joining Network algorithm (epsilon=0).

## Supplementary Material

**Supplementary Figure S1. Maximum likelihood phylogenetic trees supporting the use of *Chlorophyta* phylum as the outgroup for ABCE phylogenetic studies.** (A) Unrooted tree of 152 ABCE amino acid sequences. Representatives of *Chlorophyta* formed a separate cluster with high bootstrap support. (B) Tree with ABCE amino acid sequences from 15 selected plant species rooted on HsABCE1.

**Supplementary Figure S2. Cladogram of FeS domain of the 152 plant ABCEs.** There was a total of 113 positions in the final dataset. The tree was constructed using the Maximum Likelihood method and JTT+G model. Color coding of the selected proteins refers to the affiliation to the bigger plant phyla or families.

**Supplementary Figure S3. Cladogram of NBD1 domain of the 152 plant ABCEs.** There was a total of 249 positions in the final dataset. The tree was constructed using the Maximum Likelihood method and JTT+G model. Color coding of the selected proteins refers to the affiliation to the bigger plant phyla or families.

**Supplementary Figure S4. Cladogram of NBD2 domain of the 152 plant ABCEs.** There was a total of 293 positions in the final dataset. The tree was constructed using the Maximum Likelihood method and JTT+G model. Color coding of the selected proteins refers to the affiliation to the bigger plant phyla or families.

**Supplementary Figure S5. Images of 22 *Arabidopsis* ecotypes. White scale bar depicts 1 cm on the images.**

**Supplementary Figure S6. Alignment of human HsABCE1 and *Arabidopsis thaliana* AtABCE2 together with respective domains and conserved regions.**

**Supplementary Table S1. List of the 76 plant species used within the *ABCE* phylogenetic analysis.** Number of plant *ABCE* genes identified per species, database used, genome assembly version and genome ploidy level are depicted.

**Supplementary Table S2. Whole-genome duplications (WGDs) detected in the plant species analyzed in the study**. Data is based on the results of One Thousand Plant Transcriptomes Initiative published in 2019.

**Supplementary Table S3. List of the 152 *ABCE* genes identified from 76 plant species together with protein length and amino acid sequence similarity to AtABCE2.**

**Supplementary Table S4. Non-synonymous SNPs found in AtABCE1 and AtABCE2.**

**Supplementary Table S5. Primers used within this study.**

**Supplementary File S1. A Maximum Likelihood phylogenetic tree based on 152 plant ABCE full-length amino acid sequences.** This data was used to create Figure 4 and is in Newick file format.

**Supplementary File S2. A Maximum Likelihood phylogenetic tree based on selected 83 plant ABCE full-length amino acid sequences from the *Brassicaceae* family (32), *Poaceae* family (48) and *A. trichopoda* (2) together with *C. reinhardtii* (1) as the root of the tree.** This data was used to create Figure 5 and is in Newick file format.

**Supplementary File S3. A Maximum Likelihood phylogenetic tree based on 152 plant ABCE full-length CDS sequences.** This data was used to create Figure 6 and is in Newick file format.

## Data Availability Statement

Multiple sequence alignments of plant ABCE protein sequences used in this study have been made publicly available at Dryad repository (Jakobson, Liina (2022), Plant ABCEs, Dryad, Dataset, https://doi.org/10.5061/dryad.vhhmgqnz2).

## Acknowledgements

This work was supported by Postdoctoral grant SS458 to L. J. from Tallinn University of Technology. J.S. was funded by a Natural Sciences and Engineering Research Council of Canada Discovery Grant to Colin Garroway. The research was conducted using the equipment purchased within the framework of the Project “Plant Biology Infrastructure – TAIM” funded by the EU Regional Development Fund (2014-2020.4.01.20-0282).

