## Supplementary material for "Phylogenetic analysis of *ABCE* genes across the plant kingdom": Supplementary Figures_20230929.pdf

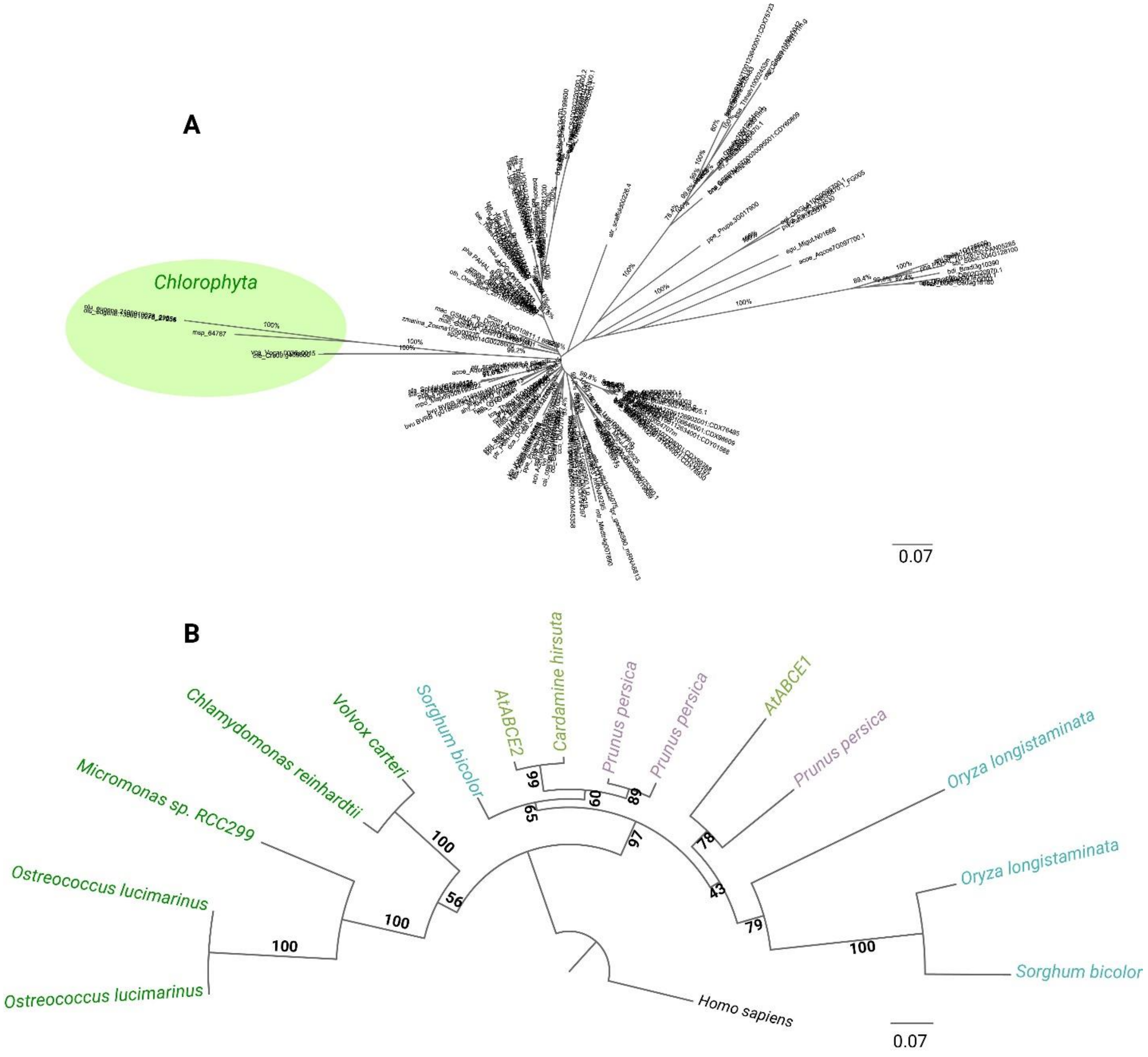


**Supplementary Figure S1. Cladograms supporting the use of Chlorophyta phylum as the outgroup for ABCE phylogenetic studies**. A) Unrooted tree of 152 ABCE amino acid sequences. Representatives of Chlorophyta formed a separate cluster with high bootstrap support. B) Cladogram with ABCE amino acid sequences from 15 selected plant species together with human HsABCE1.

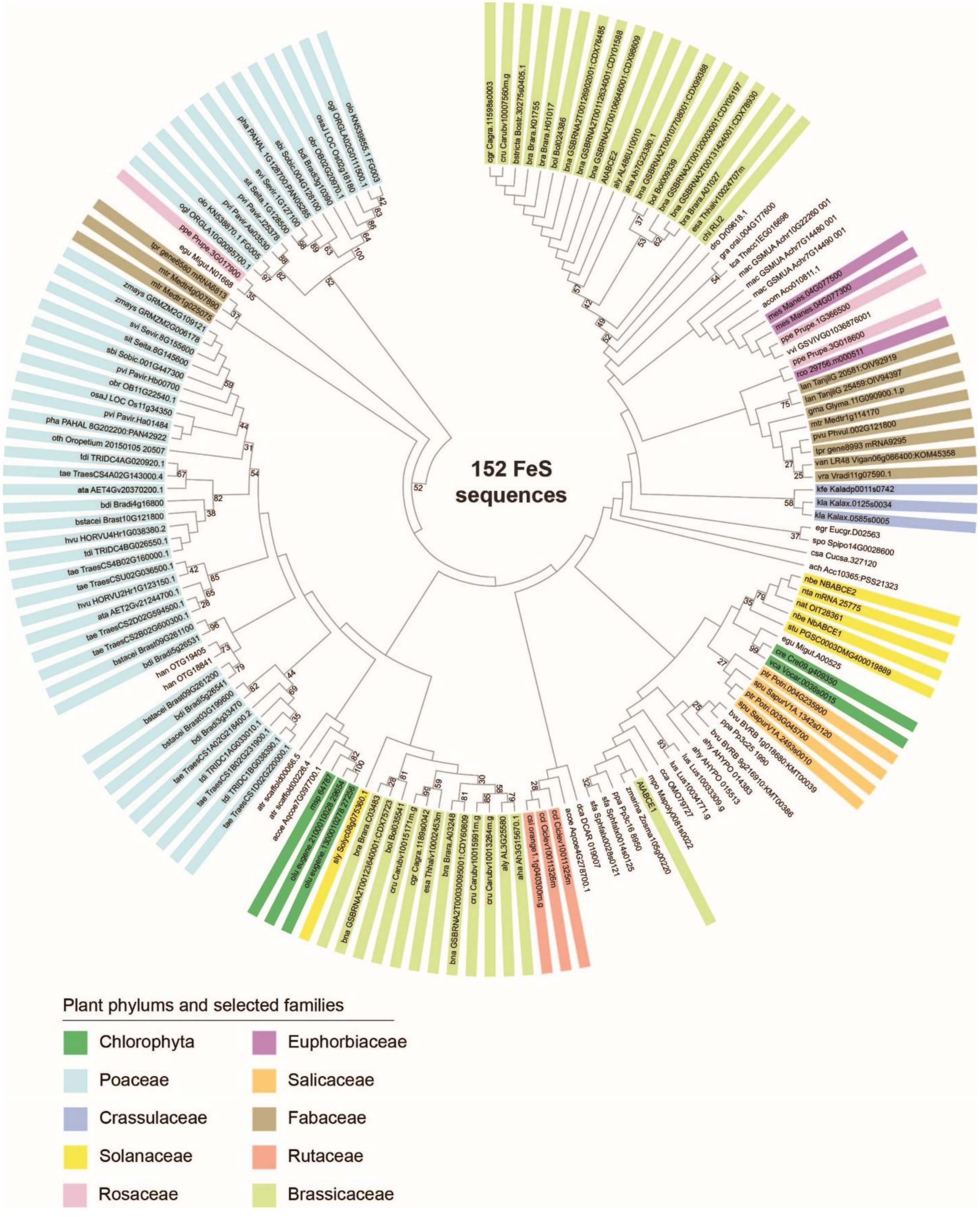


**Supplementary Figure S2. Cladogram of FeS domain of the 152 plant ABCEs.** There were a total of 113 positions in the final dataset. The tree was constructed using the Maximum Likelihood method and JTT+G model. Color coding of the selected proteins refers to the affiliation to the bigger plant phyla or families.

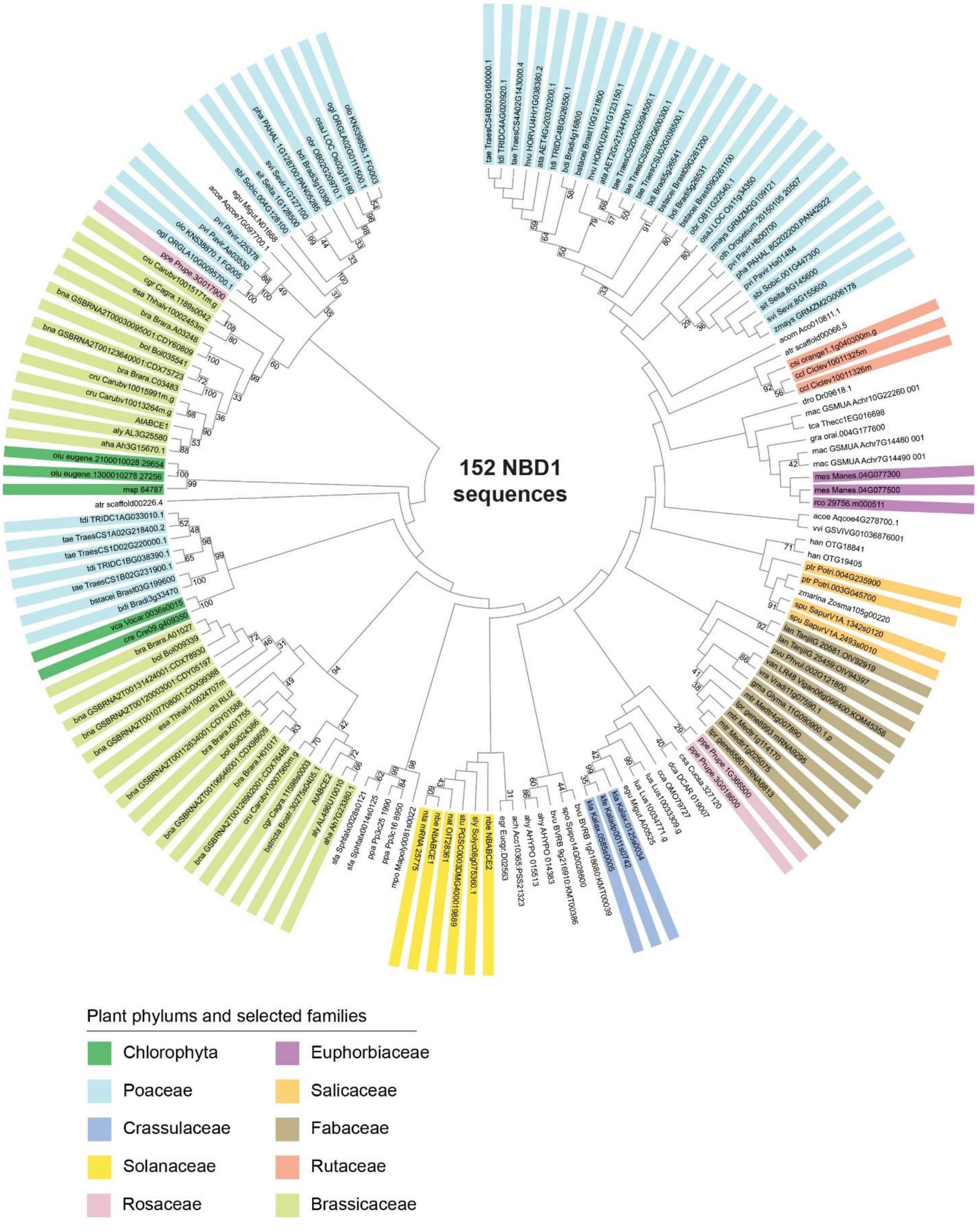


**Supplementary Figure S3. Cladogram of NBD1 domain of the 152 plant ABCEs.** There were a total of 249 positions in the final dataset. The tree was constructed using the Maximum Likelihood method and JTT+G model. Color coding of the selected proteins refers to the affiliation to the bigger plant phyla or families.

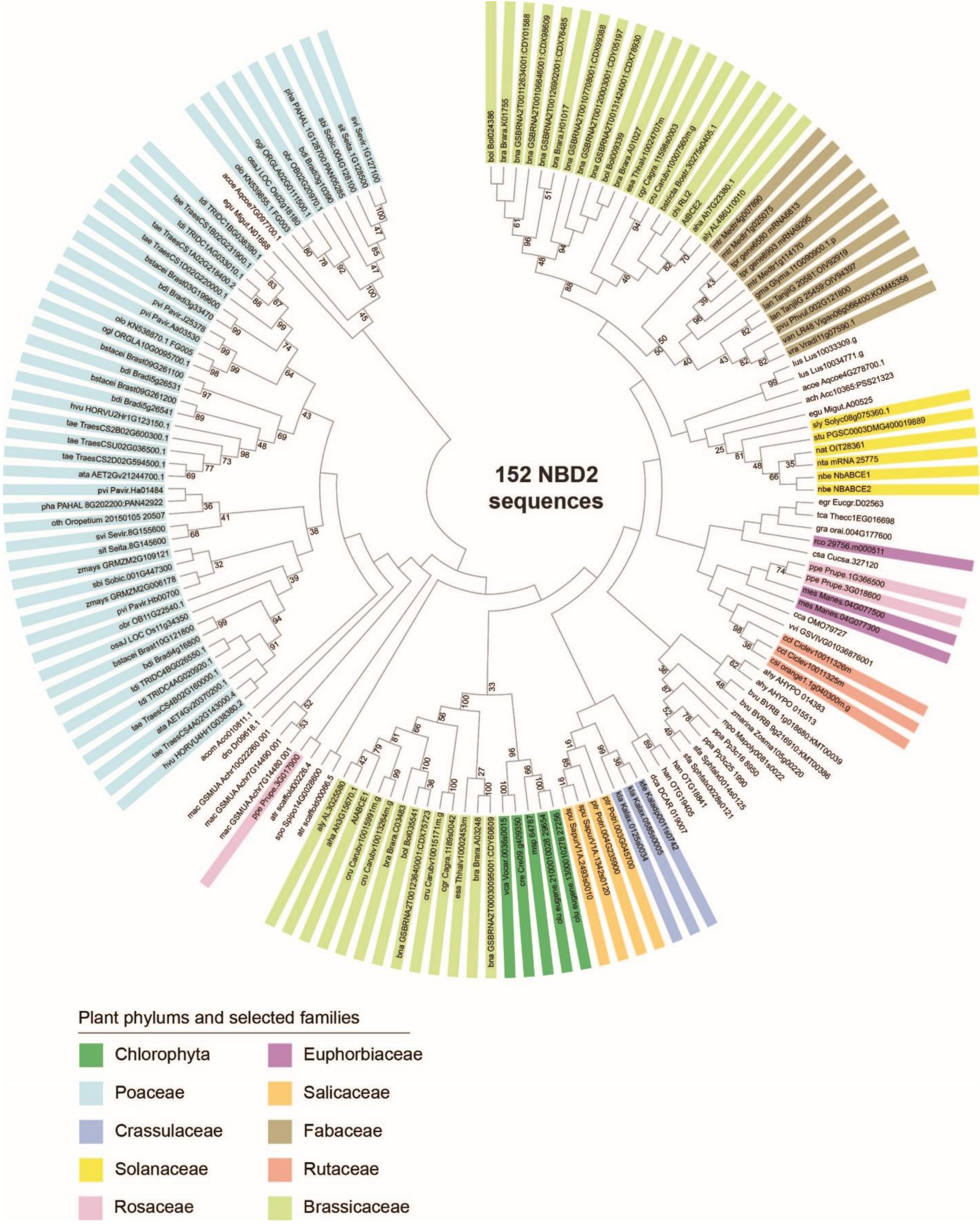


**Supplementary Figure S4. Cladogram of NBD2 domain of the 152 plant ABCEs.** There were a total of 293 positions in the final dataset. The tree was constructed using the Maximum Likelihood method and JTT+G model. Color coding of the selected proteins refers to the affiliation to the bigger plant phyla or families.

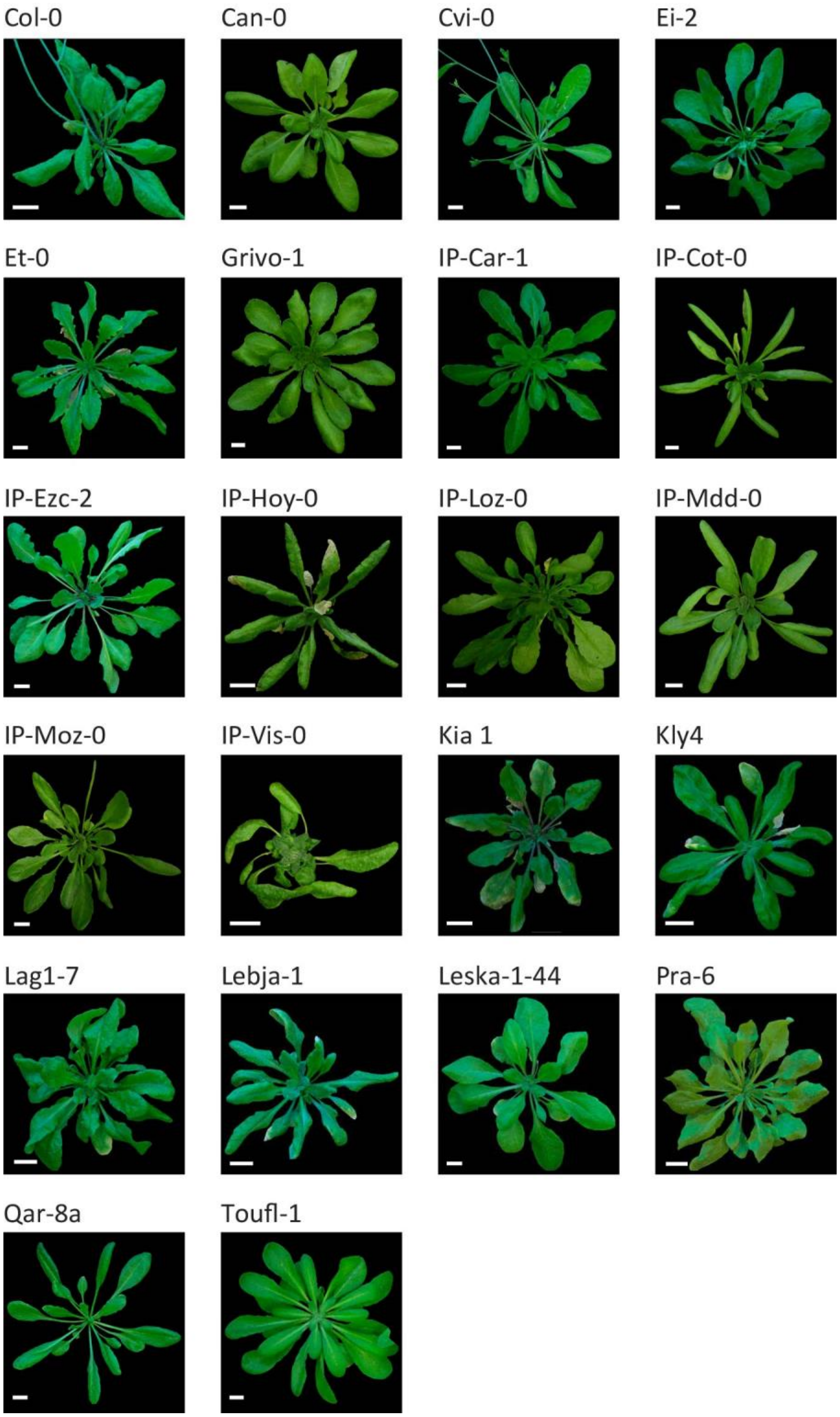


**Supplementary Figure S5. Images of 22 Arabidopsis ecotypes.** White scale bar depicts 1 cm on the images.

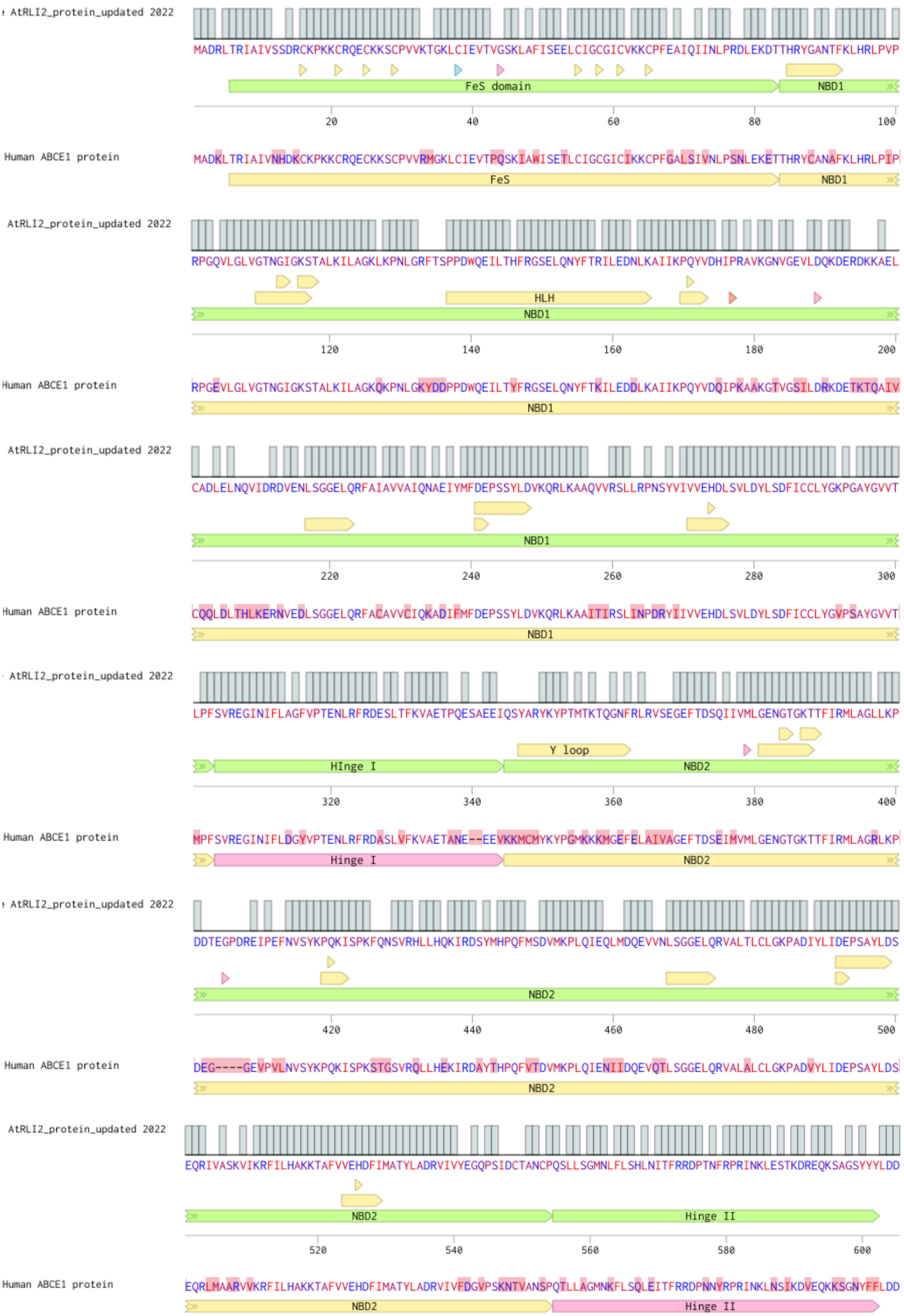


**Supplementary Figure S6. Amino acid sequence alignment of human ABCE1 vs AtABCE2.**
