## Supplementary material for "Phylogenetic analysis of *ABCE* genes across the plant kingdom": Supplementary Tables_20230929.pdf

**Supplementary Table S1. List of the 76 plant species used within the *ABCE* phylogenetic analysis.** Number of plant *ABCE* genes identified per species, database used, genome assembly version and genome ploidy level are depicted.

| No | Latin name | English name | Abbreviated name | No of genes | Ploidy | Dataset | Assembly version |
|---|---|---|---|---|---|---|---|
| 1 | *Actinidia chinensis Red5* | golden kiwifruit | Ach | 1 | diploid | Ensembl | Red5_PS1_1.69.0 |
| 2 | *Aegilops tauschii* | Tausch's goatgrass, rough-spike hard grass | Ata | 2 | diploid | Ensembl | Aet_v4.0 |
| 3 | *Amaranthus hypochondriacus* | Prince's-feather | Ahy | 2 | diploid | Phytozome | v1.0 |
| 4 | *Amborella trichopoda* | amborella | Atr | 2 | diploid endosperm?? | Phytozome | v1.0 |
| 5 | *Ananas comosus* | pineapple | Acom | 1 | diploid | Phytozome | v3 |
| 6 | *Aquilegia coerulea* | colorado blue columbine | Acoe | 2 | diploid | Phytozome | v3.1 |
| 7 | *Arabidopsis halleri* | - | Aha | 2 | diploid | Phytozome | v2.1.0 |
| 8 | *Arabidopsis lyrata* | - | Aly | 2 | diploid | Phytozome | v2.1 |
| 9 | *Arabidopsis thaliana* | thale cress | Ath | 2 | diploid | Phytozome | TAIR10 |
| 10 | *Beta vulgaris* | beet | Bvu | 2 | diploid | Ensembl | RefBeet-1.2.2 |
| 11 | *Boechera stricta* | drummond's rockcress | Bstricta | 1 | diploid | Phytozome | v1.2 |
| 12 | *Brachypodium distachyon* | purple false brome, stiff brome | Bdi | 5 | diploid | Phytozome | V3.1 |
| 13 | *Brachypodium stacei* | - | Bstacei | 4 | diploid | Phytozome | v1.1 |
| 14 | *Brassica napus* | rapeseed, oilseed rape | Bna | 8 | allopolyploidy | Ensembl | AST_PRJEB5043_v1 |
| 15 | *Brassica oleracea* | wild cabbage | Bol | 3 | diploid | Phytozome | v1.0 |
| 16 | *Brassica rapa* | field mustard | Bra | 5 | diploid | Phytozome | FPsc v1.3 |
| 17 | *Capsella grandiflora* | grand shepherd's-purse | Cgr | 2 | diploid | Phytozome | v1.1 |
| 18 | *Capsella rubella* | pink shepherd's-purse | Cru | 4 | diploid | Phytozome | v1.0 |
| 19 | *Cardamine hirsuta* | hairy bittercress | Chi | 1 | diploid | Gan et al., 2016 | MIPS_CARH_v3.8 |
| 20 | *Chlamydomonas reinhardtii* | green alga | Cre | 1 | haploid | Phytozome | v5.5 |
| 21 | *Citrus clementina* | clementine | Ccl | 2 | diploid | Phytozome | v1.0 |
| 22 | *Citrus sinensis* | sweet orange | Csi | 1 | diploid | Phytozome | v1.1 |
| 23 | *Corchorus capsularis* | white jute | Cca | 1 | diploid | Ensembl | CCACVL1_1.0 |
| 24 | *Cucumis sativus* | cucumber | Csa | 1 | diploid | Phytozome | v1.0 |
| 25 | *Daucus carota* | wild carrot, bird's nest, bishop's lace | Dca | 1 | diploid | Phytozome | v2.0 |
| 26 | *Dioscorea rotundata* | white yam, Guinea yam | Dro | 1 | diploid | Ensembl | TDr96_F1_Pseudo_Chromosome_v1.0 |
| 27 | *Erythranthe guttata* | seep monkeyflower | Egu | 2 | diploid | Phytozome | v2.0 |
| 28 | *Eucalyptus grandis* | flooded gum, rose gum | Egr | 1 | diploid | Phytozome | v2.0 |
| 29 | *Eutrema salsugineum* | saltwater cress | Esa | 2 | diploid | Phytozome | v1.0 |
| 30 | *Glycine max* | soybean | Gma | 1 | partially diploidized tetraploid | Phytozome | v2.0 |
| 31 | *Gossypium raimondii* | cotton plant endemic to Peru | Gra | 1 | diploid | Phytozome | v2.1 |
| 32 | *Helianthus annuus* | common sunflower | Han | 2 | diploid | Ensembl | HanXRQr1.0 |
| 33 | *Hordeum vulgare* | barley | Hvu | 2 | diploid | Ensembl | IBSC_v2 |
| 34 | *Kalanchoe fedtschenkoi* | kalanchoë | Kfe | 1 | diploid | Phytozome | v1.1 |
| 35 | *Kalanchoe laxiflora* | Milky Widow's Thrill | Kla | 2 | diploid | Phytozome | v1.1 |
| 36 | *Linum usitatissimum* | common flax, linseed | Lus | 2 | diploid | Phytozome | v1.0 |
| 37 | *Lupinus angustifolius* | narrow-leaved blue lupine | Lan | 2 | diploid | Ensembl | LupAngTanjil_v1.0 |
| 38 | *Manihot esculenta* | cassava, manioc, yuca | Mes | 2 | diploid | Phytozome | v6.1 |
| 39 | *Marchantia polymorpha* | common liverwort, umbrella liverwort | Mpo | 1 | haploid/diploid | Phytozome | v3.1 |
| 40 | *Medicago truncatula* | barrelclover, strong-spined medick, barrel medic | Mtr | 3 | diploid | Phytozome | Mt4.0v1 |
| 41 | *Micromonas sp. RCC299* | green alga | Msp | 1 | haploid | Phytozome | v3.0 |
| 42 | *Musa acuminata* | banana native to Southeast Asia | Mac | 3 | double haploid | Phytozome | v1 |
| 43 | *Nicotiana attenuata* | coyote tobacco | Nat | 1 | diploid | Ensembl | NIATTr2 |
| 44 | *Nicotiana benthamiana* | tobacco | Nbe | 2 | allotetraploid | Kourelis et al. 2019 | v1.0.1 |
| 45 | *Nicotiana tabacum* | cultivated tobacco | Nta | 1 | allotetraploid | solgenomics.net | TN90 protein sequences |
| 46 | *Oropetium thomaeum* | - | Oth | 1 | diploid | Phytozome | v1.0 |
| 47 | *Oryza brachyantha* | rice grass native to tropical Africa | Obr | 2 | diploid | Ensembl | v1.4b |
| 48 | *Oryza glaberrima* | african rice | Ogl | 2 | diploid | Ensembl | V1 |
| 49 | *Oryza longistaminata* | african wild rice | Olo | 2 | diploid | Ensembl | v1.0 |
| 50 | *Oryza sativa Japonica Group* | sinica rice | OsaJ | 2 | diploid | Phytozome | v7_JGI |

| No | Latin name | English name | Abbreviated name | No of genes | Ploidy | Dataset | Assembly version |
|---|---|---|---|---|---|---|---|
| 51 | *Ostreococcus lucimarinus* | green alga | Olu | 2 | haploid | Phytozome | v2.0 |
| 52 | *Panicum hallii ecotype FIL2* | hall's panicgrass | Pha | 2 | diploid | Ensembl | PHallii_v3.1 |
| 53 | *Panicum virgatum* | switchgrass | Pvi | 4 | tetraploid | Phytozome | v1.1 |
| 54 | *Phaseolus vulgaris* | common bean, French bean | Pvu | 1 | diploid | Phytozome | v2.1 |
| 55 | *Physcomitrella patens* | spreading earthmoss | Ppa | 2 | haploid/diploid | Phytozome | v3.3 |
| 56 | *Populus trichocarpa* | black cottonwood, California poplar | Ptr | 2 | diploid with haploid chromosomes | Phytozome | v3.0 |
| 57 | *Prunus persica* | peach | Ppe | 3 | diploid | Phytozome | v2.1 |
| 58 | *Ricinus communis* | castor bean | Rco | 1 | diploid | Phytozome | v0.1 |
| 59 | *Salix purpurea* | purple willow | Spu | 2 | diploid | Phytozome | v1.0 |
| 60 | *Setaria italica* | foxtail millet | Sit | 2 | diploid | Phytozome | v2.2 |
| 61 | *Setaria viridis* | green foxtail, wild foxtail millet | Svi | 2 | diploid | Phytozome | v1.1 |
| 62 | *Solanum lycopersicum* | tomato | Sly | 1 | diploid | Phytozome | iTAG2.4 |
| 63 | *Solanum tuberosum* | potato | Stu | 1 | tetraploid | Phytozome | v4.03 |
| 64 | *Sorghum bicolor* | sorghum, great millet | Sbi | 2 | diploid | Phytozome | v3.1.1 |
| 65 | *Sphagnum fallax* | flat-topped bogmoss | Sfa | 2 | haploid | Phytozome | v0.5 |
| 66 | *Spirodela polyrhiza* | common duckmeat | Spo | 1 | diploid | Phytozome | v2 |
| 67 | *Zea mays Ensembl-18* | maize, corn | Zmays | 2 | diploid | Phytozome | AGPv3 |
| 68 | *Zostera marina* | common eelgrass, seawrack | Zmarina | 1 | diploid | Phytozome | v2.2 |
| 69 | *Theobroma cacao* | cacao tree | Tca | 1 | diploid | Phytozome | v1.1 |
| 70 | *Trifolium pratense* | red clover | Tpr | 2 | diploid | Phytozome | v2 |
| 71 | *Triticum aestivum* | common wheat, bread wheat | Tae | 8 | hexaploid | Ensembl | IWGSC |
| 72 | *Triticum dicoccoides* | emmer wheat, hulled wheat | Tdi | 4 | tetraploid | Ensembl | WEWSeq_v.1.0 |
| 73 | *Vigna angularis* | adzuki bean, English red mung bean | Van | 1 | diploid | Ensembl | Vigan1.1 |
| 74 | *Vigna radiata* | mung bean | Vra | 1 | diploid | Ensembl | Vradiata_ver6 |
| 75 | *Vitis vinifera* | common grape vine | Vvi | 1 | diploid | Phytozome | Genoscope.12X |
| 76 | *Volvox carteri* | colonial green alga | Vca | 1 | haploid/diploid | Phytozome | v2.1 |

**Supplementary Table S2. Whole-genome duplications (WGDs) detected in the plant species analyzed in the study.** Data is based on the results of One Thousand Plant Transcriptomes Initiative published in 2019.

| Species | No of *ABCE* genes | No of WGDs | WGDs |
|---|---|---|---|
| *Volvox carteri* | 1 | 0 | |
| *Ostreococcus lucimarinus* | 2 | 0 | |
| *Micromonas sp. RCC299* | 1 | 0 | |
| *Marchantia polymorpha* | 1 | 0 | |
| *Chlamydomonas reinhardtii* | 1 | 1 | CHMOα |
| *Physcomitrella patens* | 2 | 1 | PHPAα |
| *Sphagnum fallax* | 2 | 1 | SPPAα |
| *Amborella trichopoda* | 2 | 2 | GINKα, AMBOα |
| *Zostera marina* | 1 | 2 | GINKα, AMBOα |
| *Brachypodium stacei* | 4 | 2 | GINKα, AMBOα |
| *Brachypodium distachyon* | 5 | 2 | GINKα, AMBOα |
| *Hordeum vulgare* | 2 | 2 | GINKα, AMBOα |
| *Aegilops tauschii* | 2 | 2 | GINKα, AMBOα |
| *Triticum dicoccoides* | 4 | 2 | GINKα, AMBOα |
| *Triticum aestivum* | 8 | 2 | GINKα, AMBOα |
| *Beta vulgaris* | 2 | 3 | GINKα, AMBOα, ARTHγ |
| *Vitis vinifera* | 1 | 3 | GINKα, AMBOα, ARTHγ |
| *Prunus persica* | 3 | 3 | GINKα, AMBOα, ARTHγ |
| *Ricinus communis* | 1 | 3 | GINKα, AMBOα, ARTHγ |
| *Citrus sinensis* | 1 | 3 | GINKα, AMBOα, ARTHγ |
| *Citrus clementina* | 2 | 3 | GINKα, AMBOα, ARTHγ |
| *Theobroma cacao* | 1 | 3 | GINKα, AMBOα, ARTHγ |
| *Helianthus annuus* | 2 | 3 | GINKα, AMBOα, ARTHγ |
| *Daucus carota* | 1 | 3 | GINKα, AMBOα, ARTHγ |
| *Cucumis sativus* | 1 | 3 | GINKα, AMBOα, ARTHγ |
| *Medicago truncatula* | 3 | 3 | GINKα, AMBOα, ARTHγ |
| *Trifolium pratense* | 2 | 3 | GINKα, AMBOα, ARTHγ |
| *Vigna radiata* | 1 | 3 | GINKα, AMBOα, ARTHγ |
| *Vigna angularis* | 1 | 3 | GINKα, AMBOα, ARTHγ |
| *Gossypium raimondii* | 1 | 3 | GINKα, AMBOα, ARTHγ |
| *Corchorus capsularis* | 1 | 3 | GINKα, AMBOα, ARTHγ |
| *Cardamine hirsuta* | 1 | 3 | GINKα, AMBOα, ARTHγ |
| *Boechera stricta* | 1 | 3 | GINKα, AMBOα, ARTHγ |
| *Eutrema salsugineum* | 2 | 3 | GINKα, AMBOα, ARTHγ |
| *Ananas comosus* | 1 | 3 | GINKα, AMBOα, ORSAγ |
| *Oropetium thomaeum* | 1 | 3 | GINKα, AMBOα, ORSAγ |
| *Aquilegia coerulea* | 2 | 3 | GINKα, AMBOα, PASOβ |
| *Spirodela polyrhiza* | 1 | 3 | GINKα, AMBOα, SPPOα |
| *Amaranthus hypochondriacus* | 2 | 4 | GINKα, AMBOα, ARTHγ, AMREα |
| *Erythranthe guttata* | 2 | 4 | GINKα, AMBOα, ARTHγ, ANMAα |
| *Kalanchoe fedtschenkoi* | 1 | 4 | GINKα, AMBOα, ARTHγ, CRAPα |
| *Kalanchoe laxiflora* | 2 | 4 | GINKα, AMBOα, ARTHγ, CRAPα |
| *Phaseolus vulgaris* | 1 | 4 | GINKα, AMBOα, ARTHγ, GLSOβ |
| *Linum usitatissimum* | 2 | 4 | GINKα, AMBOα, ARTHγ, LIUSα |
| *Solanum lycopersicum* | 1 | 4 | GINKα, AMBOα, ARTHγ, LYBAα |
| *Solanum tuberosum* | 1 | 4 | GINKα, AMBOα, ARTHγ, LYBAα |
| *Nicotiana tabacum* | 1 | 4 | GINKα, AMBOα, ARTHγ, LYBAα |
| *Nicotiana attenuata* | 1 | 4 | GINKα, AMBOα, ARTHγ, LYBAα |

| **Species** | **No of *ABCE* genes** | **No of WGDs** | **WGDs** |
|---|---|---|---|
| *Nicotiana benthamiana* | 2 | 4 | GINKα, AMBOα, ARTHγ, LYBAα |
| *Manihot esculenta* | 2 | 4 | GINKα, AMBOα, ARTHγ, MAESα |
| *Populus trichocarpa* | 2 | 4 | GINKα, AMBOα, ARTHγ, SAACα |
| *Salix purpurea* | 2 | 4 | GINKα, AMBOα, ARTHγ, SAACα |
| *Eucalyptus grandis* | 1 | 4 | GINKα, AMBOα, ARTHγ, SYMIα |
| *Dioscorea rotundata* | 1 | 4 | GINKα, AMBOα, ORSAγ, DIVIα |
| *Actinidia chinensis Red5* | 1 | 5 | GINKα, AMBOα, ARTHγ, ACCHα, ACCHβ |
| *Glycine max* | 1 | 5 | GINKα, AMBOα, ARTHγ, GLSOα, GLSOβ |
| *Lupinus angustifolius* | 2 | 5 | GINKα, AMBOα, ARTHγ, LUPOα, GLSOβ |
| *Musa acuminata* | 3 | 5 | GINKα, AMBOα, ORSAγ, MUACα, MALEα |
| *Sorghum bicolor* | 2 | 5 | GINKα, AMBOα, ORSAγ, ORSAα, ORSAβ |
| *Panicum virgatum* | 4 | 5 | GINKα, AMBOα, ORSAγ, ORSAα, ORSAβ |
| *Panicum hallii ecotype FIL2* | 2 | 5 | GINKα, AMBOα, ORSAγ, ORSAα, ORSAβ |
| *Oryza glaberrima* | 2 | 5 | GINKα, AMBOα, ORSAγ, ORSAα, ORSAβ |
| *Oryza longistaminata* | 2 | 5 | GINKα, AMBOα, ORSAγ, ORSAα, ORSAβ |
| *Oryza brachyantha* | 2 | 5 | GINKα, AMBOα, ORSAγ, ORSAα, ORSAβ |
| *Oryza sativa Japonica Group* | 2 | 5 | GINKα, AMBOα, ORSAγ, ORSAα, ORSAβ |
| *Zea mays* | 2 | 5 | GINKα, AMBOα, ORSAγ, ORSAα, ORSAβ |
| *Setaria italica* | 2 | 5 | GINKα, AMBOα, ORSAγ, ORSAα, ORSAβ |
| *Setaria viridis* | 2 | 5 | GINKα, AMBOα, ORSAγ, ORSAα, ORSAβ |
| *Brassica napus* | 8 | 6 | GINKα, AMBOα, ARTHγ, BRNIα, ARTHα, ARTHβ |
| *Brassica oleracea* | 3 | 6 | GINKα, AMBOα, ARTHγ, BRNIα, ARTHα, ARTHβ |
| *Brassica rapa* | 5 | 6 | GINKα, AMBOα, ARTHγ, BRNIα, ARTHα, ARTHβ |
| *Capsella grandiflora* | 2 | 6 | GINKα, AMBOα, ARTHγ, BRNIα, ARTHα, ARTHβ |
| *Capsella rubella* | 4 | 6 | GINKα, AMBOα, ARTHγ, BRNIα, ARTHα, ARTHβ |
| *Arabidopsis thaliana* | 2 | 6 | GINKα, AMBOα, ARTHγ, BRNIα, ARTHα, ARTHβ |
| *Arabidopsis lyrata* | 2 | 6 | GINKα, AMBOα, ARTHγ, BRNIα, ARTHα, ARTHβ |
| *Arabidopsis halleri* | 2 | 6 | GINKα, AMBOα, ARTHγ, BRNIα, ARTHα, ARTHβ |

**Supplementary Table S3. List of the 152 *ABCE* genes identified from 76 plant species together with protein length and amino acid sequence similarity to AtABCE2.**

| No | Latin species name | Species acronym in this study | Locus/gene/transcript name | Name used within this study | Peptide length | Amino acid similarity with AtABCE2 |
|---|---|---|---|---|---|---|
| 1 | *Actinidia chinensis Red5* | ach | *CEY00_Acc10365; PSS21323* | ach_Acc10365:PSS21323 | 605 | 95.9% |
| 2 | *Aegilops tauschii* | ata | *AET2Gv21244700.1* | ata_AET2Gv21244700.1 | 604 | 92.4% |
| 3 | *Aegilops tauschii* | ata | *AET4Gv20370200.1* | ata_AET4Gv20370200.1 | 604 | 93.2% |
| 4 | *Amaranthus hypochondriacus* | ahy | *AHYPO_014383-AR* | ahy_AHYPO_014383 | 605 | 95.0% |
| 5 | *Amaranthus hypochondriacus* | ahy | *AHYPO_015513-AR* | ahy_AHYPO_015513 | 605 | 95.4% |
| 6 | *Amborella trichopoda* | atr | *evm_27.model.AmTr_v1.0_scaffold00066.5* | atr_scaffold00066.5 | 606 | 95.5% |
| 7 | *Amborella trichopoda* | atr | *evm_27.model.AmTr_v1.0_scaffold00226.4* | atr_scaffold00226.4 | 605 | 89.4% |
| 8 | *Ananas comosus* | acom | *Aco010811.1* | acom_Aco010811.1 | 605 | 95.5% |
| 9 | *Aquilegia coerulea* | acoe | *Aqcoe4G278700.1* | acoe_Aqcoe4G278700.1 | 604 | 96.4% |
| 10 | *Aquilegia coerulea* | acoe | *Aqcoe7G097700.1* | acoe_Aqcoe7G097700.1 | 590 | 85.4% |
| 11 | *Arabidopsis halleri* | aha | *Ah3G15670.1* | aha_Ah3G15670.1 | 603 | 88.1% |
| 12 | *Arabidopsis halleri* | aha | *Ah7G23380.1* | aha_Ah7G23380.1 | 605 | 99.7% |
| 13 | *Arabidopsis lyrata* | aly | *AL3G25580.t1* | aly_AL3G25580 | 581 | 87.6% |
| 14 | *Arabidopsis lyrata* | aly | *AL486U10010.t1* | aly_AL486U10010 | 605 | 99.5% |
| 15 | *Arabidopsis thaliana* | ath | *AT3G13640.1; AtABCE1* | AtABCE1 | 603 | 87.1% |
| **16** | ***Arabidopsis thaliana*** | **ath** | ***AT4G19210.1; AtABCE2*** | **AtABCE2** | **605** | **100.0%** |
| 17 | *Beta vulgaris* | bvu | *BVRB_1g018680; KMT00039* | bvu_BVRB_1g018680:KMT00039 | 605 | 93.4% |
| 18 | *Beta vulgaris* | bvu | *BVRB_9g216910; KMT00386* | bvu_BVRB_9g216910:KMT00386 | 605 | 94.7% |
| 19 | *Boechera stricta* | bstricta | *Bostr.30275s0405.1* | bstricta_Bostr.30275s0405.1 | 605 | 99.0% |
| 20 | *Brachypodium distachyon* | bdi | *Bradi3g10390.1* | bdi_Bradi3g10390 | 603 | 78.6% |
| 21 | *Brachypodium distachyon* | bdi | *Bradi3g33470.1* | bdi_Bradi3g33470 | 603 | 88.3% |
| 22 | *Brachypodium distachyon* | bdi | *Bradi4g16800.1* | bdi_Bradi4g16800 | 604 | 93.9% |
| 23 | *Brachypodium distachyon* | bdi | *Bradi5g26531.1* | bdi_Bradi5g26531 | 604 | 92.7% |
| 24 | *Brachypodium distachyon* | bdi | *Bradi5g26541.1* | bdi_Bradi5g26541 | 604 | 93.4% |
| 25 | *Brachypodium stacei* | bstacei | *Brast03G199600.1* | bstacei_Brast03G199600 | 608 | 88.5% |
| 26 | *Brachypodium stacei* | bstacei | *Brast09G261100.1* | bstacei_Brast09G261100 | 604 | 93.0% |
| 27 | *Brachypodium stacei* | bstacei | *Brast09G261200.1* | bstacei_Brast09G261200 | 604 | 93.0% |
| 28 | *Brachypodium stacei* | bstacei | *Brast10G121800.1* | bstacei_Brast10G121800 | 604 | 93.9% |
| 29 | *Brassica napus* | bna | *BnaA01g37290D; CDY60809* | bna_GSBRNA2T00030095001:CDY60809 | 603 | 88.1% |
| 30 | *Brassica napus* | bna | *BnaA03g43940D; CDX98609* | bna_GSBRNA2T00106646001:CDX98609 | 605 | 99.0% |
| 31 | *Brassica napus* | bna | *BnaC01g11640D; CDX99388* | bna_GSBRNA2T00107708001:CDX99388 | 605 | 98.2% |
| 32 | *Brassica napus* | bna | *BnaC07g35780D; CDY01588* | bna_GSBRNA2T00112634001:CDY01588 | 605 | 99.0% |
| 33 | *Brassica napus* | bna | *BnaC05g46060D; CDY05197* | bna_GSBRNA2T00120003001:CDY05197 | 605 | 98.0% |
| 34 | *Brassica napus* | bna | *BnaC03g38060D; CDX75723* | bna_GSBRNA2T00123640001:CDX75723 | 603 | 85.4% |
| 35 | *Brassica napus* | bna | *BnaA08g09150D; CDX76485* | bna_GSBRNA2T00126902001:CDX76485 | 605 | 98.8% |
| 36 | *Brassica napus* | bna | *BnaA01g09970D; CDX78930* | bna_GSBRNA2T00131424001:CDX78930 | 605 | 98.0% |
| 37 | *Brassica oleracea* | bol | *Bol009339* | bol_Bol009339 | 605 | 98.2% |
| 38 | *Brassica oleracea* | bol | *Bol024386* | bol_Bol024386 | 605 | 99.0% |
| 39 | *Brassica oleracea* | bol | *Bol035541* | bol_Bol035541 | 603 | 85.4% |
| 40 | *Brassica rapa* | bra | *Brara.A01027.1* | bra_Brara.A01027 | 605 | 98.0% |
| 41 | *Brassica rapa* | bra | *Brara.A03248.1* | bra_Brara.A03248 | 603 | 88.1% |
| 42 | *Brassica rapa* | bra | *Brara.C03483.1* | bra_Brara.C03483 | 603 | 85.4% |
| 43 | *Brassica rapa* | bra | *Brara.H01017.1* | bra_Brara.H01017 | 605 | 98.8% |
| 44 | *Brassica rapa* | bra | *Brara.K01755.1* | bra_Brara.K01755 | 605 | 99.0% |
| 45 | *Capsella grandiflora* | cgr | *Cagra.11598s0003.1* | cgr_Cagra.11598s0003 | 605 | 99.2% |
| 46 | *Capsella grandiflora* | cgr | *Cagra.1189s0042.1* | cgr_Cagra.1189s0042 | 604 | 86.2% |
| 47 | *Capsella rubella* | cru | *Carubv10007560m* | cru_Carubv10007560m.g | 605 | 99.2% |
| 48 | *Capsella rubella* | cru | *Carubv10013264m* | cru_Carubv10013264m.g | 603 | 87.9% |
| 49 | *Capsella rubella* | cru | *Carubv10015171m* | cru_Carubv10015171m.g | 604 | 86.4% |
| 50 | *Capsella rubella* | cru | *Carubv10015991m* | cru_Carubv10015991m.g | 603 | 87.9% |
| 51 | *Cardamine hirsuta* | chi | *CARHR226120; ChRLI2* | chi_RLI2 | 605 | 99.3% |
| 52 | *Chlamydomonas reinhardtii* | cre | *Cre09.g409350.t1.2* | cre_Cre09.g409350 | 618 | 86.7% |
| 53 | *Citrus clementina* | ccl | *Ciclev10011325m* | ccl_Ciclev10011325m | 605 | 95.0% |
| 54 | *Citrus clementina* | ccl | *Ciclev10011326m* | ccl_Ciclev10011326m | 605 | 95.0% |
| 55 | *Citrus sinensis* | csi | *orange1.1g040300m* | csi_orange1.1g040300m.g | 605 | 94.9% |
| 56 | *Corchorus capsularis* | cca | *CCACVL1_13469; OMO79727* | cca_OMO79727 | 605 | 94.2% |
| 57 | *Cucumis sativus* | csa | *Cucsa.327120.1* | csa_Cucsa.327120 | 605 | 95.9% |
| 58 | *Daucus carota* | dca | *DCAR_019007* | dca_DCAR_019007 | 605 | 94.5% |
| 59 | *Dioscorea rotundata* | dro | *Dr09618; Dr09618.1* | dro_Dr09618.1 | 625 | 93.7% |
| 60 | *Erythranthe guttata* | egu | *Migut.A00525.1* | egu_Migut.A00525 | 605 | 95.0% |
| 61 | *Erythranthe guttata* | egu | *Migut.N01668.1* | egu_Migut.N01668 | 604 | 85.4% |
| 62 | *Eucalyptus grandis* | egr | *Eucgr.D02563.1* | egr_Eucgr.D02563 | 605 | 94.9% |
| 63 | *Eutrema salsugineum* | esa | *Thhalv10002453m* | esa_Thhalv10002453m | 599 | 86.6% |
| 64 | *Eutrema salsugineum* | esa | *Thhalv10024707m* | esa_Thhalv10024707m | 605 | 98.5% |
| 65 | *Glycine max* | gma | *Glyma.11G090900.1* | gma_Glyma.11G090900.1.p | 606 | 95.4% |
| 66 | *Gossypium raimondii* | gra | *Gorai.004G177600.2* | gra_orai.004G177600 | 605 | 96.0% |
| 67 | *Helianthus annuus* | han | *OTG18841* | han_OTG18841 | 605 | 94.7% |
| 68 | *Helianthus annuus* | han | *OTG19405* | han_OTG19405 | 605 | 94.5% |
| 69 | *Hordeum vulgare* | hvu | *HORVU2Hr1G123150.1* | hvu_HORVU2Hr1G123150.1 | 611 | 92.2% |
| 70 | *Hordeum vulgare* | hvu | *HORVU4Hr1G038380.2* | hvu_HORVU4Hr1G038380.2 | 604 | 93.4% |

| No | Latin species name | Species acronym in this study | Locus/gene/transcript name | Name used within this study | Peptide length | Amino acid similarity with AtABCE2 |
|---|---|---|---|---|---|---|
| 71 | *Kalanchoe fedtschenkoi* | kfe | *Kaladp0011s0742.1* | kfe_Kaladp0011s0742 | 605 | 94.7% |
| 72 | *Kalanchoe laxiflora* | kla | *Kalax.0125s0034.1* | kla_Kalax.0125s0034 | 605 | 94.7% |
| 73 | *Kalanchoe laxiflora* | kla | *Kalax.0585s0005.1* | kla_Kalax.0585s0005 | 605 | 94.7% |
| 74 | *Linum usitatissimum* | lus | *Lus10033309* | lus_Lus10033309.g | 605 | 94.9% |
| 75 | *Linum usitatissimum* | lus | *Lus10034771* | lus_Lus10034771.g | 605 | 94.9% |
| 76 | *Lupinus angustifolius* | lan | *TanjilG_20581; OIV92919* | lan_TanjilG_20581:OIV92919 | 606 | 94.9% |
| 77 | *Lupinus angustifolius* | lan | *TanjilG_25459; OIV94397* | lan_TanjilG_25459:OIV94397 | 606 | 94.5% |
| 78 | *Manihot esculenta* | mes | *Manes.04G077300.1* | mes_Manes.04G077300 | 605 | 95.7% |
| 79 | *Manihot esculenta* | mes | *Manes.04G077500.1* | mes_Manes.04G077500 | 605 | 95.7% |
| 80 | *Marchantia polymorpha* | mpo | *Mapoly0081s0022.2* | mpo_Mapoly0081s0022 | 605 | 94.5% |
| 81 | *Medicago truncatula* | mtr | *Medtr1g025075.1* | mtr_Medtr1g025075 | 607 | 92.6% |
| 82 | *Medicago truncatula* | mtr | *Medtr1g114170.1* | mtr_Medtr1g114170 | 606 | 94.5% |
| 83 | *Medicago truncatula* | mtr | *Medtr4g007890.1* | mtr_Medtr4g007890 | 601 | 88.5% |
| 84 | *Micromonas sp. RCC299* | msp | *64787* | msp_64787 | 614 | 85.6% |
| 85 | *Musa acuminata* | mac | *GSMUA_Achr10T22260_001* | mac_GSMUA_Achr10G22260_001 | 605 | 95.7% |
| 86 | *Musa acuminata* | mac | *GSMUA_Achr7T14480_001* | mac_GSMUA_Achr7G14480_001 | 605 | 95.5% |
| 87 | *Musa acuminata* | mac | *GSMUA_Achr7T14490_001* | mac_GSMUA_Achr7G14490_001 | 605 | 95.5% |
| 88 | *Nicotiana attenuata* | nat | *ABCE2; OIT28361* | nat_OIT28361 | 606 | 95.7% |
| 89 | *Nicotiana benthamiana* | nbe | *Niben101Scf08193g00024.1; NbABCE1* | nbe_NbABCE1 | 606 | 95.0% |
| 90 | *Nicotiana benthamiana* | nbe | *Niben101Scf02548g06001.1; NbABCE2* | nbe_NBABCE2 | 606 | 95.4% |
| 91 | *Nicotiana tabacum* | nta | *mRNA_25775* | nta_mRNA_25775 | 606 | 95.5% |
| 92 | *Oropetium thomaeum* | oth | *Oropetium_20150105_20507A* | oth_Oropetium_20150105_20507 | 604 | 95.5% |
| 93 | *Oryza brachyantha* | obr | *OB02G20970; OB02G20970.1* | obr_OB02G20970.1 | 607 | 80.1% |
| 94 | *Oryza brachyantha* | obr | *OB11G22540; OB11G22540.1* | obr_OB11G22540.1 | 604 | 95.0% |
| 95 | *Oryza glaberrima* | ogl | *ORGLA02G0111500.1* | ogl_ORGLA02G0111500.1 | 607 | 81.3% |
| 96 | *Oryza glaberrima* | ogl | *ORGLA10G0095700.1* | ogl_ORGLA10G0095700.1 | 593 | 84.1% |
| 97 | *Oryza longistaminata* | olo | *KN538870.1_FG005* | olo_KN538870.1_FG005 | 626 | 81.2% |
| 98 | *Oryza longistaminata* | olo | *KN539855.1_FG003* | olo_KN539855.1_FG003 | 676 | 81.5% |
| 99 | *Oryza sativa Japonica Group* | osaJ | *LOC_Os02g18180.1* | osaJ_LOC_Os02g18180 | 608 | 81.3% |
| 100 | *Oryza sativa Japonica Group* | osaJ | *LOC_Os11g34350.1* | osaJ_LOC_Os11g34350 | 604 | 95.0% |
| 101 | *Ostreococcus lucimarinus* | olu | *27256* | olu_eugene.1300010278_27256 | 611 | 83.1% |
| 102 | *Ostreococcus lucimarinus* | olu | *29654* | olu_eugene.2100010028_29654 | 611 | 83.1% |
| 103 | *Panicum hallii ecotype FIL2* | pha | *PAHAL_1G128700; PAN05285* | pha_PAHAL_1G128700:PAN05285 | 604 | 79.8% |
| 104 | *Panicum hallii ecotype FIL2* | pha | *PAHAL_8G202200; PAN42922* | pha_PAHAL_8G202200:PAN42922 | 604 | 95.5% |
| 105 | *Panicum virgatum* | pvi | *Pavir.Aa03530.1* | pvi_Pavir.Aa03530 | 603 | 84.0% |
| 106 | *Panicum virgatum* | pvi | *Pavir.Ha01484.1* | pvi_Pavir.Ha01484 | 604 | 95.5% |
| 107 | *Panicum virgatum* | pvi | *Pavir.Hb00700.1* | pvi_Pavir.Hb00700 | 604 | 95.5% |
| 108 | *Panicum virgatum* | pvi | *Pavir.J25378.1* | pvi_Pavir.J25378 | 603 | 84.4% |
| 109 | *Phaseolus vulgaris* | pvu | *Phvul.002G121800.1* | pvu_Phvul.002G121800 | 606 | 95.2% |
| 110 | *Physcomitrella patens* | ppa | *Pp3c16_8950V3.1* | ppa_Pp3c16_8950 | 605 | 93.9% |
| 111 | *Physcomitrella patens* | ppa | *Pp3c25_1990V3.1* | ppa_Pp3c25_1990 | 605 | 93.6% |
| 112 | *Populus trichocarpa* | ptr | *Potri.003G045700.1* | ptr_Potri.003G045700 | 611 | 94.7% |
| 113 | *Populus trichocarpa* | ptr | *Potri.004G235900.1* | ptr_Potri.004G235900 | 605 | 95.5% |
| 114 | *Prunus persica* | ppe | *Prupe.1G366500.1* | ppe_Prupe.1G366500 | 605 | 95.0% |
| 115 | *Prunus persica* | ppe | *Prupe.3G017900.1* | ppe_Prupe.3G017900 | 601 | 87.4% |
| 116 | *Prunus persica* | ppe | *Prupe.3G018600.1* | ppe_Prupe.3G018600 | 605 | 94.9% |
| 117 | *Ricinus communis* | rco | *29756.m000511* | rco_29756.m000511 | 591 | 95.4% |
| 118 | *Salix purpurea* | spu | *SapurV1A.1342s0120.1* | spu_SapurV1A.1342s0120 | 605 | 95.4% |
| 119 | *Salix purpurea* | spu | *SapurV1A.2493s0010.1* | spu_SapurV1A.2493s0010 | 605 | 95.4% |
| 120 | *Setaria italica* | sit | *Seita.1G128500.1* | sit_Seita.1G128500 | 604 | 79.1% |
| 121 | *Setaria italica* | sit | *Seita.8G145600.1* | sit_Seita.8G145600 | 604 | 95.4% |
| 122 | *Setaria viridis* | svi | *Sevir.1G127100.1* | svi_Sevir.1G127100 | 604 | 79.1% |
| 123 | *Setaria viridis* | svi | *Sevir.8G155600.1* | svi_Sevir.8G155600 | 604 | 95.4% |
| 124 | *Solanum lycopersicum* | sly | *Solyc08g075360.1.1* | sly_Solyc08g075360.1 | 606 | 93.4% |
| 125 | *Solanum tuberosum* | stu | *PGSC0003DMT400051207* | stu_PGSC0003DMG400019889 | 605 | 95.5% |
| 126 | *Sorghum bicolor* | sbi | *Sobic.001G447300.1* | sbi_Sobic.001G447300 | 604 | 95.5% |
| 127 | *Sorghum bicolor* | sbi | *Sobic.004G128100.1* | sbi_Sobic.004G128100 | 606 | 78.1% |
| 128 | *Sphagnum fallax* | sfa | *Sphfalx0014s0125.1* | sfa_Sphfalx0014s0125 | 605 | 93.6% |
| 129 | *Sphagnum fallax* | sfa | *Sphfalx0028s0121.1* | sfa_Sphfalx0028s0121 | 605 | 94.2% |
| 130 | *Spirodela polyrhiza* | spo | *Spipo14G0028600* | spo_Spipo14G0028600 | 605 | 94.7% |
| 131 | *Zea mays Ensembl-18* | zmays | *GRMZM2G006178_T01* | zmays_GRMZM2G006178 | 604 | 95.4% |
| 132 | *Zea mays Ensembl-18* | zmays | *GRMZM2G109121_T02* | zmays_GRMZM2G109121 | 604 | 95.2% |
| 133 | *Zostera marina* | zmarina | *Zosma105g00220.1* | zmarina_Zosma105g00220 | 605 | 92.4% |
| 134 | *Theobroma cacao* | tca | *Thecc1EG016698t1* | tca_Thecc1EG016698 | 605 | 95.5% |
| 135 | *Trifolium pratense* | tpr | *Tp57577_TGAC_v2_mRNA6813* | tpr_gene6580_mRNA6813 | 603 | 89.0% |
| 136 | *Trifolium pratense* | tpr | *Tp57577_TGAC_v2_mRNA9295* | tpr_gene8993_mRNA9295 | 606 | 94.0% |
| 137 | *Triticum aestivum* | tae | *TraesCS1A02G218400.2* | tae_TraesCS1A02G218400.2 | 599 | 87.6% |
| 138 | *Triticum aestivum* | tae | *TraesCS1B02G231900.1* | tae_TraesCS1B02G231900.1 | 599 | 87.8% |
| 139 | *Triticum aestivum* | tae | *TraesCS1D02G220000.1* | tae_TraesCS1D02G220000.1 | 599 | 87.6% |
| 140 | *Triticum aestivum* | tae | *TraesCS2B02G600300.1* | tae_TraesCS2B02G600300.1 | 604 | 92.5% |

| No | Latin species name | Species acronym in this study | Locus/gene/transcript name | Name used within this study | Peptide length | Amino acid similarity with AtABCE2 |
|---|---|---|---|---|---|---|
| 141 | *Triticum aestivum* | tae | *TraesCS2D02G594500.1* | tae_TraesCS2D02G594500.1 | 604 | 92.4% |
| 142 | *Triticum aestivum* | tae | *TraesCS4A02G143000.4* | tae_TraesCS4A02G143000.4 | 604 | 93.5% |
| 143 | *Triticum aestivum* | tae | *TraesCS4B02G160000.1* | tae_TraesCS4B02G160000.1 | 604 | 93.5% |
| 144 | *Triticum aestivum* | tae | *TraesCSU02G036500.1* | tae_TraesCSU02G036500.1 | 604 | 92.2% |
| 145 | *Triticum dicoccoides* | tdi | *TRIDC1AG033010.1* | tdi_TRIDC1AG033010.1 | 596 | 87.8% |
| 146 | *Triticum dicoccoides* | tdi | *TRIDC1BG038390.1* | tdi_TRIDC1BG038390.1 | 602 | 86.9% |
| 147 | *Triticum dicoccoides* | tdi | *TRIDC4AG020920.1* | tdi_TRIDC4AG020920.1 | 604 | 93.5% |
| 148 | *Triticum dicoccoides* | tdi | *TRIDC4BG026550.1* | tdi_TRIDC4BG026550.1 | 604 | 93.5% |
| 149 | *Vigna angularis* | van | *LR48_Vigan06g066400; KOM45358* | van_LR48_Vigan06g066400:KOM45358 | 606 | 95.0% |
| 150 | *Vigna radiata* | vra | *Vradi11g07590.1* | vra_Vradi11g07590.1 | 606 | 95.0% |
| 151 | *Vitis vinifera* | vvi | *GSVIVT01036876001* | vvi_GSVIVG01036876001 | 625 | 95.9% |
| 152 | *Volvox carteri* | vca | *Vocar.0036s0015.1* | vca_Vocar.0036s0015 | 619 | 86.9% |

**Supplementary Table S4. Non-synonymous SNPs found in *AtABCE1* and *AtABCE2* .**

***AtABCE2***

| position | SNP | Codon change | Ecotype | Selected ecotypes | Comment |
|---|---|---|---|---|---|
| 10502255 | G44S/Gly44Ser/c.130G>A | Ggt/Agt | 54 ecotypes: PYL-6, Brösarp-15-138, Ängsö-59-422, Mc-1, App1-14, App1-16, Fly2-1, Fly2-2, Hov1-7, Hov3-5, Kni-1, Rev-3, Stu-2, T1000, T1010, T1160, T610, T670, T800, T860, T930, TDr-2, TDr-8, Tomegap-2, Ei-2, Ull2-5, Edi-0, Gie-0, Kl-5, Kb-0, Nw-0, Oy-0, Petergof, Rome-1, Su-0, Nc-1, Algutsrum, Gul1-2, Hov4-1, Lund, Rev-1, St-0, FlyA 3, Kia 1, Ull-A-1, IP-Jim-1, IP-Moc-11, IP-Vaz-0, IP-Gud-3, IP-Mac-0, IP-Rib-1, IP-Urd-1, IP-Vas-0, Pra-6 | **Ei-2, Kia 1, Pra-6** | not conserved |
| 10503064 | D189E/Asp189Glu/c.567C>A | gaC/gaA | Ost-0 | Ost-0: not available | |
| 10503813 | M379T/Met379Thr/c.1136T>C | aTg/aCg | IP-Car-1 | **IP-Car-1** | not conserved |
| 10503980 | G405A/Gly405Ala/c.1214G>C | gGa/gCa | Can-0 | **Can-0** | not conserved |

***AtABCE1***

| position | SNP | Codon change | Ecotypes | Selected ecotypes | Comment |
|---|---|---|---|---|---|
| 4458801 | p.Lys588*/c.1762A>T | Aag/Tag | Kly-4 | **Kly-4** | stop gain mutation |
| 4458848 | p.Arg572Leu/c.1715G>T | cGg/cTg | 2 ecotypes: Aitba-1, Toufl-1 | **Toufl-1** | conserved Arg in Hinge II subdomain |
| 4458862 | p.Asn567Lys/c.1701C>A | aaC/aaA | IP-Moc-11 | IP-Moc-11: not available | |
| 4458966 | p.His561Leu/c.1682A>T | cAc/cTc | 997 ecotypes | all 21 ecotypes | not conserved |
| 4459002 | p.Ala549Gly/c.1646C>G | gCt/gGt | 6 ecotypes: Lag1-2, Lag1-5, Lag1-6, Lag1-7, Lag2-4, Qar-8a | **Qar-8a, Lag1-7** | substitution for similar AA residue |
| 4459219 | p.Thr477Ala/c.1429A>G | Act/Gct | Uod-7 | | not conserved |
| 4459291 | p.Leu453Phe/c.1357C>T | Ctt/Ttt | 7 ecotypes: IP-Cem-0, IP-Cor-0, IP-Fun-0, IP-Hum-2, IP-Nac-0, IP-Pun-0, IP-Ven-0 | | substitution for similar AA residue |
| 4459327 | p.Ala441Thr/c.1321G>A | Gca/Aca | 28 ecotypes: Hovdala-2, Hovdala-2, Cvi-0, Kz-9, Ak-1, Kn-0, Sei-0, Zu-1, Lag1-2, Lag1-5, Lag1-6, Lag1-7, Lag2-4, Lag2-7, Lag2-10, Bak-5, EkS 2, EkS 3, Kolyv-3, Goced-1, Podvi-1, Stara-1, Leska-1-44, Koren-1, Malak-1, Epidauros-1, Qar-8a, IP-Mdd-0, Bak-2 | **IP-Mdd-0, Cvi-0, Leska-1-44, Qar-8a, Lag1-7** | not conserved |
| 4459351 | p.His433Asp/c.1297C>G | Cat/Gat | IP-Cat-0 | | not conserved |
| 4459411 | p.Val413Leu/c.1237G>C | Gtg/Ctg | RUM-20 | | substitution for similar AA residue |
| 4459438 | p.Val404Ile/c.1210G>A | Gta/Ata | Zdarec3 | | substitution for similar AA residue |
| 4459443 | p.Glu402Val/c.1205A>T | gAg/gTg | Faneronemi-3 | | not conserved |
| 4459450 | p.Arg400Ser/c.1198C>A | Cgt/Agt | 2 ecotypes: Kz-9, Kolyv-3 | | not conserved |
| 4459450 | p.Arg400Cys/c.1198C>T | Cgt/Tgt | IP-Ses-0 | | not conserved |
| 4459453 | p.Pro399Thr/c.1195C>A | Cca/Aca | Grivo-1 | **Grivo-1** | substitution of conserved Pro in NBD2 |
| 4459607 | p.Asp373His/c.1117G>C | Gac/Cac | 6 ecotypes: Lag1-2, Lag1-5, Lag1-6, Lag1-7, Lag2-4, Qar-8a | **Lag1-7, Qar-8a** | substitution of conserved Asp in NBD2 |
| 4459652 | p.Gln358Lys/c.1072C>A | Caa/Aaa | IP-Ldd-0, Furni-1, Iasi-1, Bolin-1 | | not conserved |
| 4459687 | p.Ser346Phe/c.1037C>T | tCc/tTc | App1-14 | | not conserved |
| 4459714 | p.Thr337Arg/c.1010C>G | aCa/aGa | 21 ecotypes: Leb-3, Karag-2, Basta-1, Basta-2, Basta-3, Chaba-2, Lebja-1, Lebja-2, Masl-1, Nosov-1, Noveg-1, Noveg-2, Noveg-3, Panke-1, Rakit-1, Rakit-2, Rakit-3, Sever-1, Stepn-2, Stepn-1, Kidr-1 | **Lebja-1** | not conserved |
| 4459823 | p.Ser328Phe/c.983C>T | tCc/tTc | Cimin-1 | | not conserved |
| 4459839 | p.Arg323Cys/c.967C>T | Cgt/Tgt | IP-Moz-0 | **IP-Moz-0** | conserved Arg in Hinge I subdomain |
| 4459928 | p.Pro293Gln/c.878C>A | cCa/cAa | IP-Vis-0 | **IP-Vis-0** | substitution of conserved Pro in NBD1 |
| 4459995 | p.Val271Ile/c.811G>A | Gtt/Att | Cvi-0 | **Cvi-0** | substitution for similar AA residue |
| 4460286 | p.Leu206Phe/c.618G>T | ttG/ttT | App1-12 | | substitution of conserved Leu in NBD1 |
| 4460360 | p.Gly182Ser/c.544G>A | Ggt/Agt | 8 ecotypes: Jl-3, IP-All-0, IP-Vin-0, IP-Bra-0, IP-Cot-0, IP-Vas-0, Lecho-1, Dobra-1 | **IP-Cot-0** | substitution of conserved Gly in NBD1 |

| position | SNP | Codon change | Ecotype | Selected ecotypes | Comment |
|---|---|---|---|---|---|
| 4460508 | p.Val160Ile/c.478G>A | Gta/Ata | 58 ecotypes: Brösarp-11-135, App1-16, Boo2-3, Fly2-1, Fly2-2, Kni-1, T1010, T1160, T610, T670, T710, T780, T800, T850, T880, T900, T930, T960, T970, T980, Ga-0, LL-0, Ts-5, Wt-5, Et-0, Fr-2, Hs-0, Mnz-0, Ob-0, Old-1, Or-0, Rsch-4, Sf-1, Kent, Bå1-2, Bla-1, Lund, FlyA 3, HolA-1 2, Ull-A-1, IP-Ang-0, IP-Coc-1, IP-Elb-0, IP-Gua-1, IP-Hor-0, IP-Moc-11, IP-Mon-5, IP-Rds-0, IP-Tam-0, IP-Aru-0, IP-Cas-0, IP-Mac-0, IP-Mie-1, IP-Sal-0, IP-Tri-0, ESP-1-11, ARGE-1-15, Qui-0 | **Et-0** | substitution for similar AA residue |
| 4460511 | p.Val159Leu/c.475G>C | Gta/Cta | UKSE06-639 | | substitution for similar AA residue |
| 4460513 | p.Arg158Gln/c.473G>A | cGa/cAa | Et-0 | **Et-0** | not conserved |
| 4460571 | p.Asp139Asn/c.415G>A | Gac/Aac | IP-Mdd-0 | **IP-Mdd-0** | not conserved |
| 4460697 | p.Pro129Gln/c.386C>A | cCa/cAa | 15 ecotypes: Ak-1, Kn-0, Sei-0, Zu-1, Lag1-2, Lag1-5, Lag1-6, Lag1-7, Lag2-4, Podvi-1, Stara-1, Leska-1-44, Koren-1, Qar-8a, IP-Mdd-0 | **Lag1-7, Qar-8a,IP-Mdd-0,Leska1-44** | substitution of conserved Pro in NBD1 |
| 4460709 | p.Gly125Glu/c.374G>A | gGa/gAa | 35 ecotypes: Dör-10, Dra-3, Eden-1, Eden-5, Eden-6, Eden-7, GrÃ¶n-5, Nyl-2, Nyl-7, TÃ„L 03, TEDEN 02, TFÃ„ 04, TFÃ„ 06, TFÃ„ 07, TFÃ„ 08, TGR 01, TNY 04, FÃ¤b-2, FÃ¤b-4, Tamm-2, Tamm-27, Sanna-2, FÃ¤L 1, GrÃ¶n 12, Nyl 13, IP-Hoy-0, IP-Ria-0, IP-Are-0, IP-Boa-0, IP-Coy-0, IP-Elp-0, IP-Leg-0, IP-Loz-0, IP-Pad-0, IP-Pva-1 | **IP-Hoy-0, IP-Loz-0** | substitution of conserved Gly in NBD1 |
| 4460719 | p.Ile122Leu/c.364A>C | Atc/Ctc | IP-Adc-5 | | substitution for similar AA residue |
| 4460908 | p.Arg86Gln/c.257G>A | cGa/cAa | 19 ecotypes: Draha2, DraII-6, DraIV 1-8, DraIV 2-9, DraIV 6-22, Duk, UduI 4-9, ZdrI 2-21, Dra-0, Jm-0, Kyoto, Petergof, Da(1)-12, DraIII-1, Rak-2, IP-Bos-0, IP-Ezc-2, Galdo-1, Monte-1 | **IP-Ezc-2** | substitution of conserved Arg in Y-loop I |
| 4460921 | p.Asp82Tyr/c.244G>T | Gat/Tat | Spro 1 | | not conserved |
| 4461106 | p.Ile50Leu/c.148A>C | Ata/Cta | 2 ecotypes: Sr:3, Strand-1 | | substitution for similar AA residue |

**Table S5. Primers used within this study.**

| Primer name | Primer sequence | Comments |
|---|---|---|
| **RLI1_F4** | 5’ - GAAGGAATCAACGTGTTCTTGG - 3’ | For sequencing *AtABCE1* SNPs |
| **RLI1_R4** | 5’ - CCATCTCTCTCTGACCGATC - 3’ | For sequencing *AtABCE1* SNPs |
| **RLI1_F3** | 5’ - TTGCAGGCTTCCAATTCCAC - 3’ | For sequencing *AtABCE1* SNPs |
| **RLI1_R3** | 5’ - CTGAAGGTCAGGGATTCATCC - 3’ | For sequencing *AtABCE1* SNPs |
| **RLI1_s** | 5’ - TTGACTTGTGTATCTTGTAT - 3’ | For sequencing *AtABCE1* SNPs |
| **RLI1_R2** | 5’ - GCTGGAAAACTCAAACCAAATC - 3’ | For sequencing *AtABCE1* SNPs |
| **RLI2_F3** | 5’ - CAGAGTCCTCCAGACTGGCAAG - 3’ | For sequencing *AtABCE2* SNPs |
| **RLI2_R3** | 5’ - CCTTGAAGGTCAAAGATTCATCTC - 3’ | For sequencing *AtABCE2* SNPs |
| **RLI2_RP** | 5’ - CTGTAGGAACAAATCCAGCCA - 3’ | For sequencing *AtABCE2* SNPs |
| **RLI2_LP** | 5’ - TTCTTGGTCTGAAATTGGTGG - 3’ | For sequencing *AtABCE2* SNPs |
| **RLI2_F1** | 5’ - TGTTATTTCGTTCCTTTGCCTT - 3’ | For sequencing *AtABCE2* SNPs |
| **RLI2_as** | 5’ - CATCTTGGATATCGGAAAGAGC - 3’ | For sequencing *AtABCE2* SNPs |
